# Genomic context-dependent histone H3K36 methylation by three *Drosophila* methyltransferases and implications for dedicated chromatin readers

**DOI:** 10.1101/2024.02.06.577191

**Authors:** Muhunden Jayakrishnan, Magdalena Havlová, Václav Veverka, Catherine Regnard, Peter B. Becker

**Affiliations:** Biomedical Center, Molecular Biology Division, Ludwig-Maximilians-Universität, Munich, Germany; Institute of Organic Chemistry and Biochemistry (IOCB) of the Czech Academy of Sciences, Prague, Czech Republic; Department of Cell Biology, Faculty of Science, Charles University, Prague, Czech Republic

## Abstract

Methylation of histone H3 at lysine 36 (H3K36me3) marks active chromatin. The mark is interpreted by epigenetic readers that assist transcription and safeguard the integrity of the chromatin fiber.

The chromodomain protein MSL3 binds H3K36me3 to target X-chromosomal genes in male *Drosophila* for dosage compensation. The PWWP-domain protein JASPer recruits the JIL1 kinase to active chromatin on all chromosomes. Unexpectedly, depletion of K36me3 had variable, locus-specific effects on the interactions of those readers. This observation motivated a systematic and comprehensive study of K36 methylation in a defined cellular model.

Contrasting prevailing models, we found that K36me1, K36me2 and K36me3 each represent independent chromatin states. A gene-centric view of the changing K36 methylation landscape upon depletion of the three methyltransferases Set2, NSD and Ash1 revealed local, context-specific methylation signatures. Set2 catalyzes K36me3 predominantly at transcriptionally active euchromatin. NSD places K36me2/3 at defined loci within pericentric heterochromatin and on weakly transcribed euchromatic genes. Ash1 deposits K36me1 at regions with enhancer signatures.

The genome-wide mapping of MSL3 and JASPer suggested that they bind K36me2 in addition to K36me3, which was confirmed by direct affinity measurement. This dual specificity attracts the readers to a broader range of chromosomal locations and increases the robustness of their actions.

## Introduction

Histone modifications highlight functional chromatin states. Particular chemical modifications inform about whether large chromatin domains are permissive or repressive to transcription and demarcate regulatory sequences within these domains. Combinations of modifications, most prominently acetylation and methylation of specific histone lysines, constitute epigenomic signatures that allow interpreting eukaryotic genomes (1–3).

A prominent example of a histone modification that is associated with a specific function is the trimethylation of lysine 36 of histone H3 (H3K36me3; K36me3 for short), which maps to bodies of transcribed genes. This mark correlates to active transcription in all eukaryotes from yeast to mammals. This methylation signals the ‘transcribed’ state of chromatin to localize a wide variety of processes that go along with active transcription, such as splicing (4,5), nucleosome turnover (6) deacetylation (7,8), DNA repair (9–11), RNA methylation (12), establishment of DNA methylation (13,14) and counteraction of facultative heterochromatin spreading (15,16), as well as regulation of S-adenosyl methionine flux (17,18). For example, K36me3 recruits enzymes to reinstate the integrity of the nucleosome fiber during transcription and to repress cryptic, intragenic promoters in yeast (19). In human cells, H3K36me3 can recruit DNA methyltransferases and decreased H3K36me3 is associated with increased cryptic transcription during aging (20). In *Drosophila*, K36me3 serves as an epigenetic mark that targets dosage compensation to X-linked genes in males (21).

The effects of histone methylation are usually mediated by ‘reader’ proteins, i.e., proteins that contain domains that recognize and bind specifically to methylated lysines. As subunits of larger enzyme complexes, these reader proteins tether epigenetic regulators to specific chromatin regions (22). Reader domains for H3K36 methylation include PWWP (23,24), chromo (25,26) and tudor domains (27). We are interested in two *Drosophila* proteins for which K36me3 seems to be the main determinant for genomic distribution: MSL3, the reader subunit of the male-specific-lethal dosage compensation dosage compensation complex (DCC) that boosts the transcription on the X chromosome in male flies (28) and JASPer, the targeting subunit of the JJ-complex responsible for interphase H3S10 phosphorylation which is also enriched on the male X-chromosome (29).

Lysines may be modified by one, two or three methyl groups. Whereas K36me3 has been extensively studied, K36me1/2 is less understood. The prevailing view, mainly based on characterization of the distribution of the H3K36 methylation marks in yeast, was that K36me3 corresponded to the true functional state, while K36me1/2 were mostly methylation intermediates.

Histone methylation profiles are established by dedicated histone methyltransferases (HMTs). In mammals, H3K36 HMTs are to some extent redundant. SetD2 catalyzes K36me1/2/3, whereas NSD1, NSD2 and ASH1L catalyze K36me2 and NSD3 catalyzes K36me1 (17). In *Drosophila*, only three evolutionary conserved enzymes methylate K36, namely Set2 (alias dHypb), NSD (alias dMes4) and Ash1. Set2 (orthologous to mammalian SetD2) is an interactor of the elongating RNA polymerase II and is thought to catalyze all K36me3 (28,30). NSD (orthologous to mammalian NSD1/2/3) has been reported to localize to both eu- and heterochromatin and is known predominantly as a mono- and dimethyltransferase (30,31). Finally, Ash1 (orthologous to mammalian Ash1L) is described as dimethyltransferase whose role is developmentally restricted to a few hundred genomic loci to counteract Polycomb repression (32–34). According to the prevailing model of the division of labor, NSD first ‘premodifies’ chromatin with K36me1/2, which then serves as a substrate for Set2 to co-transcriptionally generate K36me3, establishing a linear relationship between the two enzymes (30,35,36). Ash1 is believed to function independently of the Set2-NSD tandem (34,37).

This simple view is currently being challenged by several independent observations. In *Drosophila*, K36me3, me2 and me1 are differently distributed, implying independent functions for each methylation state (2,38). Furthermore, the transcriptional and developmental consequences of NSD deletion are distinct from that of Set2 deficiency (38,39). For instance, Set2-null flies are early embryonic lethal, while NSD-null flies are viable and fertile (albeit with minor defects), arguing against an exclusive linear relationship of the two enzymes towards generating the trimethylated state. Interpretation of these organismal phenotypes is complicated by suspected cell type-specific and developmental effects.

To more systematically evaluate the function of the three *Drosophila* HMTs and to characterize the different patterns of the K36 methylation states at high resolution, we resorted to a defined tissue culture model. We mapped the individual K36 methylation states at high resolution using an enhanced ChIP protocol that combines the resolution of Micrococcal Nuclease digestion with solubilization of heterochromatin by mild shearing (29). We describe the consequences of systematic HMT depletion, individual and in combination, on the K36 methylation landscape and highlight differences in local HMT dependencies. In contrast to the prevailing model, we find that the three enzymes establish distinct methylation states at defined genomic regions.

We also monitored the distribution of the two K36me3 reader proteins MSL3 and JASPer upon HMT depletion and showed that reduction of K36me3 did not lead to a proportional reduction in reader’s binding. We propose that the robustness of the reader’s recruitment is at least in part due to their ability to bind both K36me2 and K36me3 combined with threshold effects and influence of the chromatin context.

Our systematic study uncovers an unexpected complexity of the three different H3K36 methylation states, which likely have different functions depending on the local chromatin context.

## Materials and Methods

### Cell culture and RNAi

S2-DRSC (DGRC stock #), Kc167 (DGRC stock #) cells were cultured in Schneider’s *Drosophila* Medium (Thermo Fisher), supplemented with 10% heat-inactivated Fetal Bovine Serum (FBS, Sigma-Aldrich), 100 units/mL penicillin and 0.1 mg/mL streptomycin (Sigma-Aldrich) at 26°C. RNAi against target genes was performed either in large scale (for ChIP-seq) or small scale (Western Blots, Immunofluorescence staining, RNA extraction):

#### Large scale

1×10^6^ S2 cells were seeded per well of a 6-well dish (Cellstar) (two wells per RNAi condition). Cells were washed once in serum-free medium followed by treatment with 10 µg dsRNA/well/RNAi target in 1ml serum-free medium. Two different dsRNAs were used for each target to improve RNAi efficiency. Cells were incubated for 10 min at room temperature (RT) with slight agitation and further 50 min at 26°C. Two volumes of complete growth medium were added and cells were incubated for 5 days at 26°C. At day 5, cells were resuspended and counted. 2×10^7^ cells per RNAi condition were transferred to a 75 cm^2^ flask for a second round of dsRNA treatment (80 µg dsRNA/flask/RNAi target) in 8 ml serum-free media and incubated as mentioned above. At day 10, cells were counted and processed for ChIP-seq. For Ash1, to improve knockdown efficiency, three rounds of knockdowns were performed for 12 days as described by (40).

#### Small scale

3×10^5^ S2 cells were seeded per well of a 12-well dish (Cellstar). Once adhered, cells were washed once in serum-free medium followed by treatment with 3-4 µg dsRNA/well/RNAi target in 300 µl serum-free medium. Cells were incubated for 10 min at RT with slight agitation and further 50 min at 26°C. Two volumes of complete growth medium were added and cells were incubated for 5 days at 26°C. At day 5, cells were resuspended and counted. 1.5×10^6^ cells per RNAi condition were transferred to a 6-well dish for a second round of dsRNA treatment (10 µg dsRNA/flask/RNAi target) in 1 ml serum free media and incubated as mentioned above. At day 10, cells were counted and processed for Western Blot or Immunofluorescence.

dsRNA was generated by *in vitro* transcription (NEB T7 E2050s transcription kit) from PCR products generated by the following forward and reverse primers (separated by comma):

- *gst* RNAi: TTAATACGACTCACTATAGGGAGAATGTCCCCTATACTAGGTTA, TTAATACGACTCACTATAGGGAGAACGCATCCAGGCACATTG;

The sequence of *Schistosoma japonicum* GST was amplified from pGEX-4T-1 (GE Healthcare)

- *gfp* RNAi: TTAATACGACTCACTATAGGGTGCTCAGGTAGTGGTTGTCG, TTAATACGACTCACTATAGGGCCTGAAGTTCATCTGCACCA;
- Set2_#1 RNAi: TTAATACGACTCACTATA*G*GGAGAAAATCCTTGATTCCAAGCAA, TTAATACGACTCACTATAGGGAGAAGTGGTTTCTACATTTTCGT;
- Set2_#2 RNAi: TTAATACGACTCACTATAGGGAGACACGGCTTGAGATTGCTACA, TTAATACGACTCACTATAGGGAGACATGGACATGCTTTTGTTGG;
- NSD_#1 RNAi: TTAATACGACTCACTATAGGGAGACGCGAATTCCTGAGCACGGACGCGCACTC, TTAATACGACTCACTATAGGGAGACGCTCTAGATGGACACACGCTGTTGTTGCTGTTT;
- NSD_#2 RNAi: TTAATACGACTCACTATAGGGAGACCCTCCTCTGTGAGCATCGA, TTAATACGACTCACTATAGGGAGAACAACGTTTTCGTACGTCTGG;
- Ash1_#1 RNAi: TTAATACGACTCACTATAGGGAGACTTTGTGGCCAGGACCAATCAA, TTAATACGACTCACTATAGGGAGACAGGCAAGGGATCGTGCTCGGT;
- Ash1_#2 RNAi: CTAATACGACTCACTATAGGGAGGCAGTGCCATGGAGACCC, CTAATACGACTCACTATAGGGAGCAACACCCAGCAGCGTCC

*D. virilis* 79f7Dv3 cells (29) were cultured in Schneider’s *Drosophila* Medium (Thermo Fisher), supplemented with 5% heat-inactivated FBS (Sigma-Aldrich), 100 units/mL penicillin, and 0.1 mg/mL streptomycin (Sigma-Aldrich) at 26°C.

### Immunofluorescence microscopy and analysis

Immunofluorescence microscopy analysis of S2 cells was performed as described in (41). Briefly, cells after RNAi were seeded on poly-L-lysine (Sigma #P8920, 0.01% (w/v) final concentration) coated coverslips and allowed to adhere for 1h. Coverslips were then fixed, permeabilized, blocked and incubated in primary antibodies overnight at appropriate dilutions (see table). Secondary antibody staining was performed the next day after which coverslips were incubated in DAPI and sealed onto glass slides with Vectashield mounting reagent.

Confocal images were acquired on a Leica TCS SP8 with a 63x/1.4NA oil-immersion objective. Image stacks were recorded at 100 Hz scan speed with a pixel size of 350 nm (or 50 nm for Fig S1B) and z-step size of 300 nm. Pinhole was set to 1 AU (580 nm reference wavelength). Fluorescence signals were recorded sequentially to avoid channel crosstalk. Further image processing and maximum intensity projections were done in Fiji (42). CellProfiler (43) was used to quantify histone modification immunostaining signals within DAPI defined regions (pipeline provided) and further plotted on R.

### Western Blot

2-3×10^6^ RNAi-treated cells were pelleted and lysed in 1x Laemmli Buffer at a concentration of 25,000 cells/µl. Samples were denatured 95°C for 10 mins. 6-8 µL of lysate per sample was electrophoresed on SDS ServaGel TGPrimer (14% gel for histone modifications, 8% gel for other proteins) for 1.5-2 h at 180 V. Proteins were transferred to AmershamTM ProtranTM 0.45 µM Nitrocellulose Blotting Membrane for 1.5 h at 300-400 mA in either regular transfer buffer (20% MeOH, 25 mM Tris, 192 mM Glycine) or high molecular weight transfer buffer for methyltransferases (10% MeOH, 0.037% SDS, 25 mM Tris, 192 mM Glycine). Membranes were blocked with 3% BSA for 1h at RT. The membrane was incubated with primary antibody (see table) overnight at 4°C in 3% BSA PBS washed thrice with PBS-T (1x Phosphate-Buffered Saline, 0.1% Tween-20) and incubated with secondary antibody in PBS-T for 1 h in RT. Images were acquired and quantified using the LICOR Odyssey CLx.

### RNA isolation, cDNA synthesis and RT-qPCR

Total RNA was extracted from 1.5×10^6^ cells using the RNeasy Mini kit (Qiagen) according to the manufacturer’s instructions. cDNA was synthesized from 500 ng of total RNA using Superscript III First Strand Synthesis System (Invitrogen, Cat. No 18080–051, random hexamer priming) and following standard protocol. cDNA was diluted 1:100, qPCR reaction was assembled using Fast SYBR Green Mastermix (Applied Biosystem, Cat. No 4385612) and ran on a Lightcycler 480 II (Roche) instrument. Primer efficiencies were calculated via serial dilutions. Primer sequences for Ash1 and 7sk (control) were obtained from (34) and (44), respectively.

### Chromatin Immunoprecipitation after MNase treatment (MNase ChIP-seq) with spike-in

ChIP-seq on MNase-digested chromatin and sonicated chromatin was performed as previously described (29). For spike-in ChIP-seq on MNase-digested chromatin in combination with mild sonication, S2 cells (~1.2 × 10^8^ cells) after RNAi were harvested and cross-linked for 8 min by adding 1.1 mL 10× fixing solution (50 mM HEPES pH 8.0, 100 mM NaCl, 1 mM EDTA, 0.5 mM EGTA and 10% (w/v) methanol-free formaldehyde) per 10 mL culture at RT. The reaction was stopped by adding glycine at 125 mM final concentration and incubating for 10 min on ice. Cells were washed twice in PBS before subsequent steps. For nuclei isolation, S2 cells were spiked with 5% (relative cell number) 79f7Dv3 fixed cells (processed as described for S2 cells), resuspended in PBS supplemented with 0.5% (v/v) Triton X-100 and cOmplete EDTA-free Protease Inhibitor Cocktail (Roche) (PI), volume was adjusted to 7×10^7^ cells/mL and cells incubated for 15 min at 4°C with end-over-end rotation. Nuclei were collected by centrifuging at 4°C for 10 min at 2000×*g* and washed once in PBS. For chromatin fragmentation, nuclei were spun down at 4°C for 10 min at 2000×*g*, resuspended in RIPA (10 mM Tris/HCl pH 8.0, 140 mM NaCl, 1 mM EDTA, 1% (v/v) Triton-X 100, 0.1%(v/v) SDS, 0.1% (v/v) Sodium deoxycholate) supplemented with PI and 2 mM CaCl_2_ at 7×10^7^ cells/mL. These lysates were digested in 1 mL aliquots by adding 0.6 U MNase (Sigma Aldrich), resuspended in EX-50 (50 mM KCl, 10 mM HEPES pH 7.6, 1.5 mM MgCl_2_, 0.5 mM EGTA, 10% glycerol) at 0.6 U/µL, and incubated at 37°C for 35 min with slight agitation. The reaction was stopped by adding 10 mM EGTA and placing on ice. Digested chromatin was mildly sheared further with Covaris AFA S220 using 12 × 12 tubes at 50 W peak incident power, 20% duty factor and 200 cycles per burst for 8 min at 5°C. Subsequent steps were performed as described in (29). Libraries were prepared with NEBNext Ultra II DNA Library Prep Kit for Illumina (NEB, E7645) and analyzed with either 2100 Bioanalyzer or TapeStation systems (Agilent). Libraries were sequenced on NextSeq1000 (Illumina) instrument yielding typically 20–25 million 150 bp paired-end reads per sample.

### CUT&RUN

CUT&RUN was performed as previously described (45) with the Rb α-H3K36me3 antibody diluted 1/500.

### Library preparation

For ChIP-seq samples, libraries were prepared using NEBNext Ultra II DNA Library with a starting ChIP DNA amount of 3-6 ng according to manufacturer’s instructions. All libraries were sequenced on an Illumina NextSeq1000 sequencer at the Laboratory of Functional Genomic Analysis (LAFUGA, Gene Center Munich, LMU). About 20 million paired-end reads were sequenced per sample for each of the ChIP samples.

### Antibodies

| Antibody | Western | IF | ChIP | Supplier/Reference |
| --- | --- | --- | --- | --- |
| Rb α-H3K36me1 | 1:5000 | 1:800 | 2 μL | Abcam-ab9048 |
| Rb α-H3K36me2 | 1:5000 | 1:800 | 2 μL | Abcam-ab9049 |
| Rb α-H3K36me3 | 1:5000 | 1:800 | 2 μL | Abcam-ab9050 |
| Rb α-H4K16ac | | | 2 μL | Millipore 07-329 |
| Ms α-H3K9me2 | | | 2 μL | Abcam-ab1220 |
| Rb α-H2AV | 1:500 | | | Börner and Becker, 2016 (46) |
| Gp α-MSL3 | | | 3 μL | Albig et al., 2019 (29) |
| Gp α-JASPer | | | 3 μL | Albig et al., 2019 (29) |
| Rb α-NSD | 1:1000 | | 5 μL | Bell et al., 2007 (30) |
| Ms α-NSD | 1:50 | 1:10 | | Bell et al., 2007 (30) |
| Ms α-Set2 | 1:50 | | | Bell et al., 2007 (30) |
| RBP1-S2ph (ePol) | | | 2 μL | Abcam-ab5095 |
| Lamin | 1:1000 |  |  | Gift from H Saumweber |

### Read processing

Sequence reads were Demultiplexed by JE demultiplexer (47) using the barcodes from the Illumina Index read files. Demultiplexed files were aligned to either *D. melanogaster* reference genome (BDGP6, release 104) or independently to *D. virilis* genome (droVir3. Feb 2006) using Bowtie2 (48) version 2.28.0 (parameter “--end-to-end --very-sensitive --no-unal --no-mixed --no-discordant -X 500 -I 10”) and filtered for quality using samtools 1.6 (49) with a MAPQ score cutoff of -q 10. For transposons, a custom genome containing repetitive regions was used as before (29) at the alignment step. Tag directories and input-normalized coverage files were generated using *Homer* (50) with the parameter -totalReads set to the number of reads mapped to *D. virilis* genome for spike-in normalization. Input-normalized, scaled *D. melanogaster* coverage per base pair files were visualized using the Integrative Genomics Viewer (51). Replicate coverages were first analyzed independently to confirm similarities in HMT-dependency patterns after which they were averaged for subsequent analyses. Resizing of coverages to fixed window sizes of mean signal was performed with bedops and bedmap (52).

### Peak calling and annotation

Broad domains of modified H3K36me1/2/3 were called using MACS2 v2.1.2 (53) bdgpeakcall function using parameters -l 1000 -c 3. Manipulation (filtering, merging etc.) of peak sets was performed with BEDTOOLS2 v2.28.0. Genomic annotation of peaks was done using HOMER annotatePeaks.pl script.

### Data analysis and plotting

Data analysis was conducted in R (54)(R Core Team, 2020) using tidyverse libraries. ChromoMaps were generated using R chromoMap package (55). Clustered heatmaps were made using R package ‘ComplexHeatmap’ (56). Chromatin State Annotations were derived from modENCODE (2). Gene annotations were obtained from FlyBase GTF annotations BDGP6 release 104. Only genes associated with a unique FlyBase (FBgn) ID (N=17.8k) were used for subsequent analyses.

### Differential binding analysis

To identify significant differential K36me1/2/3 regions across RNAi conditions, csaw v1.24.3 (57) was used. Reads for each unique experimental sample were counted into sliding windows of 250 bp across the entire genome. Windows were filtered using filterWindowsControl function using the corresponding Input profiles as controls. Data were binned into larger 10-kbp bins and subsequently used to calculate normalization factors using normFactors function. Differential binding was assessed for using the quasi-likelihood (QL) framework in the edgeR package v3.32.1 with robust=TRUE for glmQLFit. The design matrix was constructed using a layout specifying the RNAi treatment as well as the experimental batch. Proximal tested windows were merged into regions of maximum 3 kbp by clusterWindows with a cluster level FDR target of 0.05.

### Z-score transformation

Z-score data transformation for average signal within 5-kbp genomic windows (Figure 3) or genes (Figure 6) was performed similar to previously described method (58). Average feature coverage instead of RPKM was used in calculations. A consensus peak set representing all genomic regions containing atleast one K36 modification in any RNAi condition was further used to filter windows before Z-score represention in Figure 3.

### External Datasets

| Dataset | Source |
| --- | --- |
| Ash1 ChIP (S2 cells) | Huang et al., 2017 (34) |
| H3K36me2 ChIP in Ash1 RNAi (S2 cells) | Huang et al., 2017 (34) |
| Chromatin States | Kharchenko et al., 2011 (2) |
| H3K36me3 modENCODE (S2 cells) | modENCODE (GSE20785) |
| H3K36me3 Native-ChIP (Fly) | Chaouch et al., 2021 (58) |
| H3K36me3 modified MNase ChIP (S2 cells) | Albig et al., 2019 (29) |
| Su(var)3-7 | modENCODE (GSE23487) |
| H3K9me2 | modENCODE(GSE20792) |
| H4K20me1 | modENCODE(GSE27743) |
| H3K27ac | modENCODE(GSE20779) |
| H3K4me1 | modENCODE(GSE32826) |
| Nurf | modENCODE(GSE20829) |
| H2B-ub | modENCODE(GSE20773) |
| H3K79me3 | modENCODE(GSE45090) |
| Kdm4A | modENCODE(GSE32839) |
| Kdm2 | modENCODE(GSE45061) |
| MRG15 | modENCODE(GSE25367) |
| Beaf-32 | modENCODE(GSE20760) |
| SMC3 | modENCODE(GSE45054) |
| CTCF | modENCODE(GSE32750) |
| H3K27me3 | modENCODE(GSE20781) |
| H3.3 | Henikoff et al., 2009 (59) |
| SNR1 | Hendy et al., 2022 (60) |
| ISWI | Hendy et al., 2022 (60) |
| WT S2 cells RNAseq | Albig et al., 2019 (29) |
| Fly HMT KO RNAseq | Lindehell et al., 2021 (38) |
| FlyAtlas Tissue Expression Data | Chintapalli et al., 2007 (61) |
| H3K36me2/3 in HMT KO (C3H10T cells) | Weinberg et al., 2019 (13) |
| RNAseq of WT C3H10T cells | Weinberg et al., 2019 (13) |
| H3K36me3 in HMT dTAG cells (mESC) | Sun et al., 2023 (62) |

Genomic coordinates of datasets previously aligned to genome version dm3 were transformed to newer dm6 using liftOver tool (63).

### Recombinant protein expression and purification

JASPer full length sequence was cloned into pET derived T7 driven vector (ampicillin resistance) behind His_6_ affinity tag and TEV (Tobacco Etch Virus) protease cleavage site. After the cleavage, a cloning artefact of five amino acids (SNAAS) remained at the N-terminus of JASPer.

Chemical competent NiCo21 pRARE2 (DE3) *E. coli* (NEB C2529H with additional plasmid coding rare codons and Chloramphenicol resistance) were transformed by heat shock with JASPer plasmid and cultivated on LB agar medium (100 μg/ml ampicillin, 25 µg/ml chloramphenicol and 1% glucose) overnight in 37°C. Colonies were resuspended in LB broth (Sigma) supplemented with 100 μg/ml ampicillin, 25 μg/ml chloramphenicol and 0.5% glycerol to OD_600_= 0.05. Bacterial cultures were cultivated in LEX Epiphyte 3 bioreactor at 37°C until OD_600_ =0.8 and then transferred to 18°C. Protein expression was induced with 400 μM IPTG (isopropyl-β-D-thiogalactopyranoside). The cells were harvested after 20 h expression by centrifugation (5000 g, 4°C, 20 min), were stored in −80°C.

Bacterial cells were lysated by Emulsiflex C3 in lysis buffer (25 mM Tris-HCl pH 7.5, 1 M NaCl, 2 mM β-mercaptoethanol, 10 μM EDTA) with protease inhibitor cocktail tablet (EDTA free) and DNAse I. The lysate was cleared by centrifugation (25000 g, 4°C, 30 min) and loaded on Ni-NTA-resin (Sigma). The bound protein (with His_6_ tag) was then eluted by imidazole buffer (25 mM Tris-HCl pH 7.5, 1 M NaCl, 2 mM β-mercaptoethanol, 250 mM imidazole). The purest fractions were dialyzed in presence of TEV protease to lysis buffer overnight. JASPer without His_6_ tag was present in the flow-through fraction in the second Ni affinity chromatography. JASPer sample was concentrated using an Amicon Ultra centrifugal filtration unit and loaded onto size exclusion chromatography Superdex 200 10/300 GL in buffer 25 mM Tris-HCl pH 7.5, 150 mM NaCl, 1 mM TCEP, 10 μM EDTA). The final concentration of the purified protein was determined on a NanoDrop spectrophotometer, and its purity was evaluated by SDS-PAGE stained by Coomassie Brilliant Blue.

Nucleosomes were prepared as described in (24).

### MicroScale Thermophoresis (MST)

JASPer protein (1-475) was dialysed overnight to MST buffer without Tween (20 mM HEPES pH 7.5, 50 mM NaCl, 1 mM TCEP). The stock of exchanged protein was divided in to equal aliquots and stored in −80°C. After thawing, each aliquot was centrifuged (20000 g, 4°C, 10 min), and the protein concentration was re-measured. JASPer at constant concentration 0.4 μM was mixed in 16-step dilution series (2:1 ratio, 0.5 nM and reaching up to 16 μM) with reconstituted nucleosomes (147 bp H3K36me0/2/3). The samples were adjusted to final volume of 30 μl MST buffer (20 mM HEPES pH 7.5, 50 mM NaCl, 1 mM TCEP, 0.05% Tween-20). All reaction tubes were centrifuged and incubated for 15 minutes at room temperature. Mixtures were loaded into Monolith NT.LabelFree premium capillaries and measured at 20% LED Power and 40% MST Power on a NanoTemper Monolith NT.LabelFree device at 24°C. Each mixture was prepared twice, where each replicate was measured in technical triplicates. Values were fitted in the GraphPad Prism using a built-in equation for specific binding with Hill slope.

## Results

### H3K36 methylation states and methyltransferases mark distinct chromatin types in S2 cells

To study the genomic distribution of the histone H3K36 methylation states (and their corresponding HMTs, we generated high-resolution MNase chromatin immunoprecipitation (ChIP-seq) profiles in male *Drosophila* S2 cells. Our optimized protocol includes the solubilization of chromatin through mild shearing, yielding significantly more signals at mappable heterochromatin (Supplementary Figure S1A).

We mapped K36me1/2/3 using highly specific antibodies (64), the HMT NSD and the elongating RNA polymerase II (ePol) as a proxy for Set2 (for which no ChIP-grade antibodies exist). A recently published Ash1 ChIP profile (34) was included. We also mapped two *bona fide* K36me3 binders: JASPer, a PWWP-domain reader that colocalizes with K36me3 genome-wide and MSL3, a K36me3 chromo-domain reader restricted to the X chromosome (29). The browser views (Figure 1A) reveal the distinct distributions of considered features, in particular for K36me1/2 and 3. To visualize the differences in the distribution of the K36 methylation states as well as the associated factors at a chromosome-wide level, we generated chromoMaps representing average ChIP signals in 10-kbp bins superimposed on the chromosomes (Figure 1B, Supplementary Figure S1B). Tracks depicting annotated genes as well as K9me2 peaks highlighting the gene-poor mappable pericentric heterochromatic regions (PCH) were included for reference. Of note, the PCH region is not prominently represented on the X chromosome because it is poorly mapped in the dm6 reference genome. K36me1 is largely euchromatic, and only partly overlaps with K36me2/3. K36me2 is heterochromatic, but also present in stretches of euchromatin. Surprisingly, K36me3 is distributed rather uniformly between euchromatin and PCH, where it is not restricted to genes (Supplementary Figure S1A). Set2 (inferred from ePol) and Ash1 are largely enriched at euchromatin, while NSD showed a clear heterochromatin enrichment but is also found in euchromatic regions. The two K36me3 reader proteins JASPer and MSL3 correlate best with K36me3 as already described (Figure 1A). Of note, the chromoMaps of the X chromosome for MSL3 and JASPer predominantly illustrate the X-chromosomal enrichment due to the association to dosage compensation.

**Figure 1:**
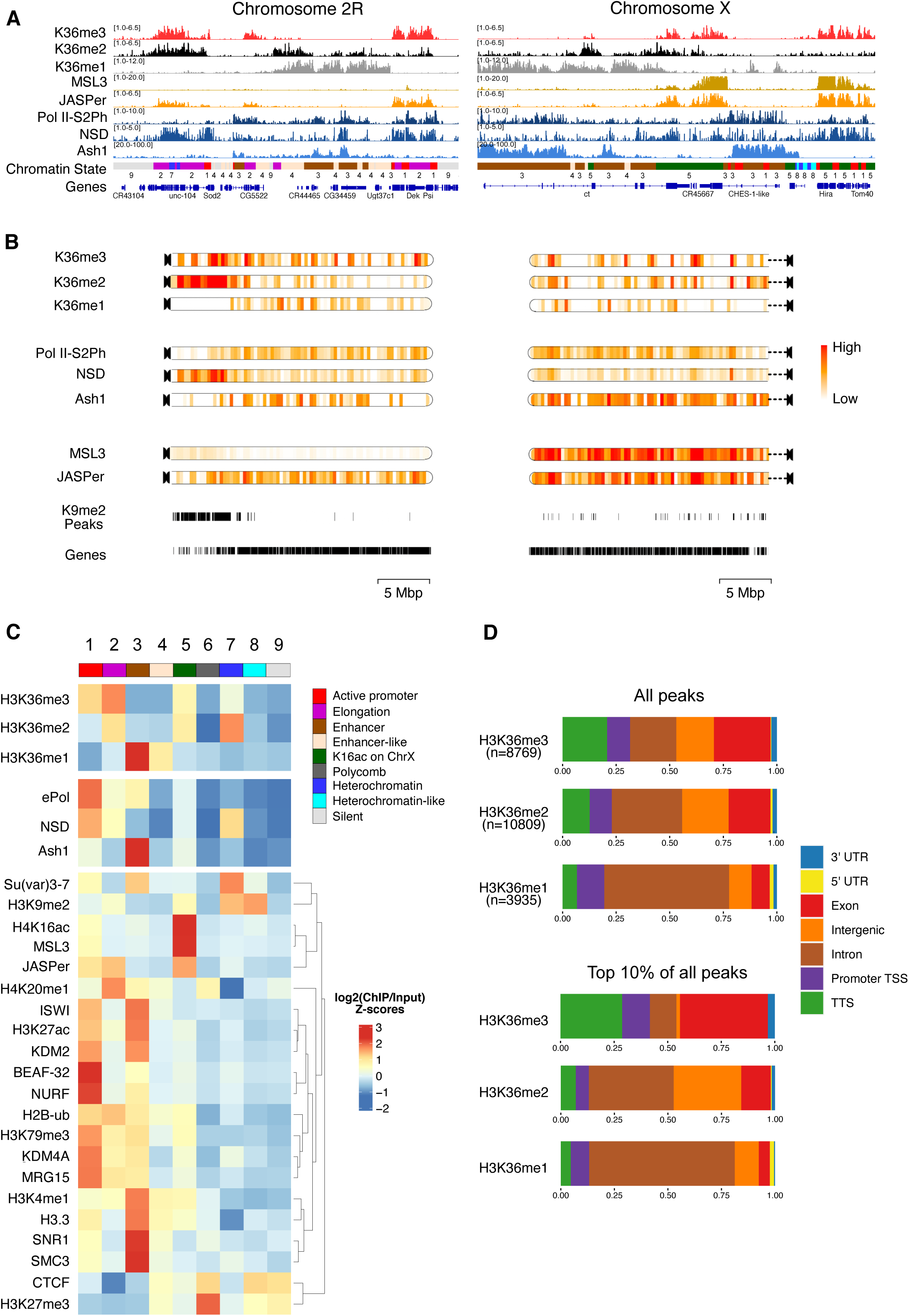
H3K36 modifications and HMTs mark distinct chromatin states in male *Drosophila* cells. A) Genome browser profiles for representative *Drosophila* chromosomes 2R and X of spike-normalized MNase-ChIP of K36me1/2/3, the K36me3 reader proteins JASPer and MSL3 and the HMTs NSD and Ash1. ‘ePol’ refers to the signal generated by an antibody against RBP1-S2ph, as a proxy for RNA-polymerase-S2ph-interacting Set2. The Ash1 profile was taken from (34). The 9-state ChromHMM (modENCODE) is color-coded and explained in (C). B) ChromoMaps representing steady-state enrichment of K36me1/2/3, HMTs and K36me3 readers in 10-kbp genomic bins for chromosomes 2R and X. Scale bars are different for each chromoMap to facilitate visualization of changes. The individual scale values for each state and chromosome are shown in Supplementary Figure S1. Tracks representing H3K9me2 peaks and annotated genes serve as a reference for mappable pericentric heterochromatin domains and transcribed chromatin, respectively. C) Chromatin State enrichment (9-state ChromHMM, (2)) for K36me1/2/3, HMTs and K36me3 readers. Published ChIP-seq/CUT&RUN profiles of histone modifications and other chromatin proteins were hierarchically clustered to highlight differences between chromatin states. D) Genomic features marked by either all K36me1/2/3 domains (top) or filtered for the strongest 10% (bottom).

We called peaks to estimate the proportion of genome decorated by each K36me mark. K36me2 and K36me3 occupy a similar proportion of the genome (~20%) while K36me1 is concentrated more local-ly (~12%). Correlating the coverage to estimated absolute modification numbers derived from late-stage embryos (65) suggests that K36me1 is the most densely placed mark, followed by K36me2 and then K36me3 (Supplementary Figure S1C).

To examine the distribution of the marks and the associated factors at a higher resolution, we next correlated ChIP enrichments to each of the 9 functional Chromatin States as defined by the modENCODE consortium (Figure 1C). We included various modENCODE ChIP datasets as well as more recently published profiles generated from published data to highlight features distinguishing the 9 States. This revealed several interesting patterns confirming the chromoMap observations. K36me3 is strongly enriched within euchromatic States 1 and 2 representing active promoters and transcribed chromatin, but also shows a mild heterochromatin enrichment (State 7). K36me2 marks similar states but is much more abundant at heterochromatin. In contrast, K36me1 is almost exclusively present at enhancer-like Chromatin States 3 and 4.

Correlating the HMT occupancies to the different K36 methylation states showed that Ash1 is most strongly enriched with K36me1 at enhancers (State 3). NSD is distributed between eu- and heterochromatin (States 1 and 7). Set2 (ePol) is rather restricted to euchromatin at transcribed regions (State 1 and 2) and enhancers (State 3) but rather not correlating with the mild K36me3 enrichment in heterochromatin (State 7). The dual distribution of NSD was confirmed by immunofluorescence confocal microscopy (Supplementary Figure S1D). In addition to a diffuse signal in euchromatin, intense speckles adjacent to (but not overlapping) DAPI-dense chromocenters were observed. Both populations were sensitive to RNA interference against NSD, confirming the specificity of the antibody.

We then intersected the peaks of K36me1/2/3 with genomic features considering the strongest peaks (top 10 percent) to highlight differences (Figure 1D). K36me1 is strongly enriched in introns, in agreement with its localization to enhancers. Strong K36me3 signals accumulate at TTS and exons, whereas K36me2 marks mostly genic features in particular introns but also some intergenic regions. In conclusion, different H3K36 methylation states label distinct types of chromatin and genomic features. NSD stands out due to its association with both eu- and heterochromatin.

### Contribution of methyltransferases to different H3K36 methylation states

To determine the contribution of HMTs to the three K36 methylation states, we depleted each HMT individually as well as in combination by RNA interference (RNAi) and followed the levels of the HMTs and K36me1/2/3 by quantitative Western blotting. The depletion of Set2 and NSD was efficient (>90% and ~80%, respectively, Supplementary Figure S2A). Because the Ash1 antibody performed poorly in Western blotting, we estimated Ash1 mRNA levels by RT-qPCR too, which revealed a 75% reduction upon RNAi (Supplementary Figure S2B).

We first followed bulk changes in K36 methylation upon HMT RNAi (Figure 2A, representative blots in Supplementary Figure S2C). Loss of Set2 led to ~75% reduction of K36me3, while K36me2 appeared unaffected. Depletion of NSD strongly reduced K36me2 (~80%), but led to only modest reduction of K36me3 (~40%). These observations are at odds with the idea that NSD only generates intermediates for Set2-dependent trimethylation. We also observed similar trends in female Kc cells, ruling out cell- or sex-specific effects (Supplementary Figure S2D, E).

**Figure 2.**
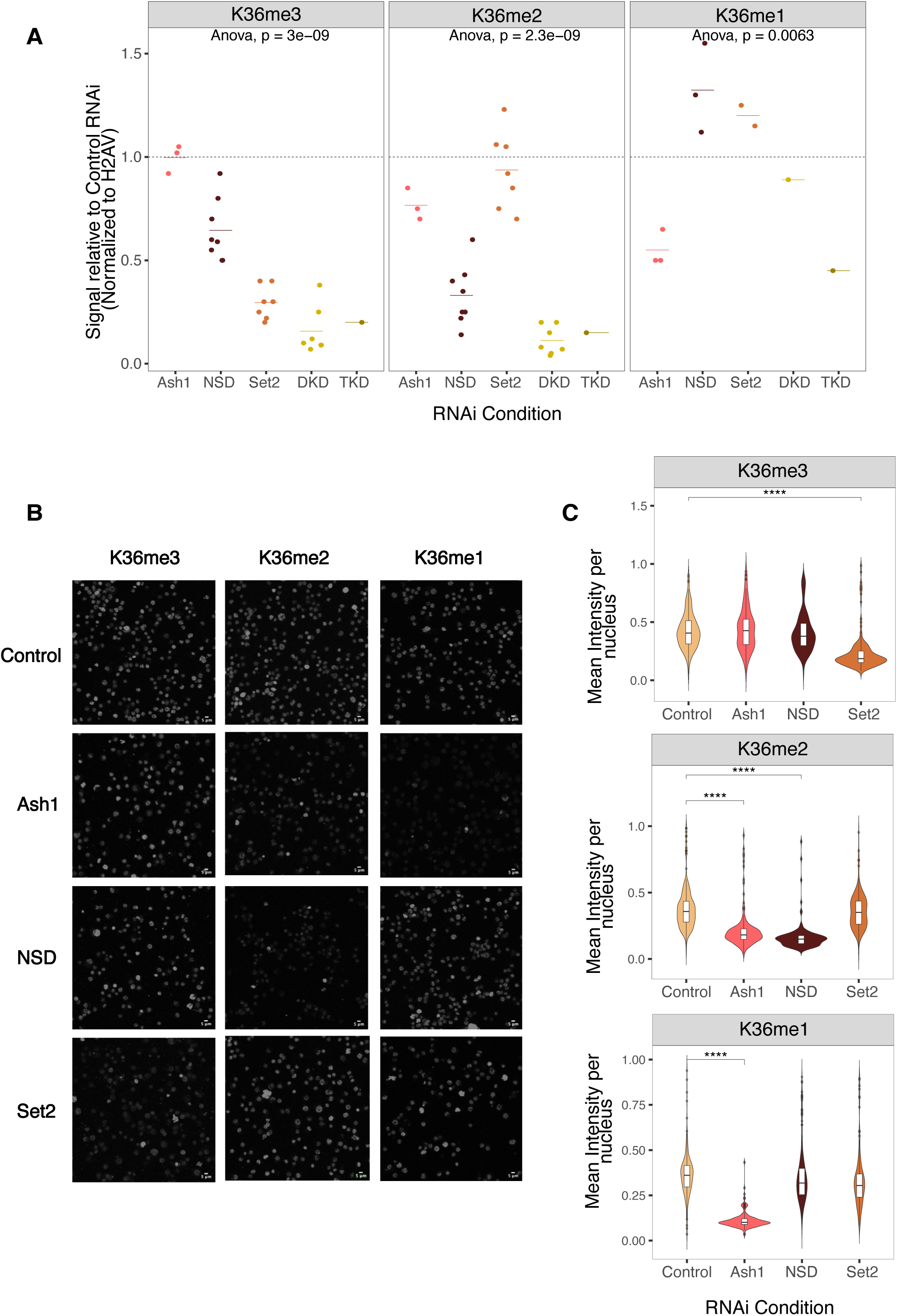
Effect of HMT depletion on H3K36 methylation bulk levels reveals distinct HMT dependencies in male cells. A) K36me1/2/3 levels in whole cell extracts from S2 cells were determined by quantitative Western blotting using specific antibodies. Cells were treated with RNAi against Ash1, NSD, Set2, NSD+Set2 (DKD) or NSD+Set2+Ash1(TKD). An irrelevant GST RNAi served as control. Representative blots are shown in Supplementary Figure S2C; Source data for all blots can be found in Zenodo Repository (see Data Availability). Values were normalized to histone H2AV signals on the same membrane and are represented as fraction relative to GST RNAi, which was run on same blot. Each dot represents an independent biological replicate. Calculated ANOVA p-values (null hypothesis: difference between means=0) are presented for each antibody. B) Representative immunofluorescence microscopy (IFM) images for α-H3K36me1/2/3 in S2 cells treated with RNAi against GST (Control), Ash1, NSD or Set2. The scale bar is 5 µm. C) Quantification of IFM images (n=~500 nuclei from 1^st^ biological replicate, shown in Figure 2B). ANOVA followed by post-hoc Tukey HSD was performed to identify groups with significantly different mean relative to GST RNAi (**** denotes a significance value of p < 0.001).

Interestingly, K36me1 was rather unaffected in RNAi of Set2 and/or NSD, arguing against the idea that NSD is the major K36me1 methyltransferase. Partial depletion of Ash1 did not affect K36me3 levels, moderately decreased K36me2 by about 20%, but led to very prominent reduction of K36me1 (50%). This reduction may be an underestimation, since the previously mentioned modENCODE study (64) reported that certain lots of K36me1 antibody have mild reactivity towards unmodified K36 in Western blot assays.

We confirmed the Western blot trends by immunofluorescence microscopy (Figure 2B), quantifying the K36me1/2/3 signals upon HMT depletion (Figure 2C, Supplementary Figure S2F). K36me1 was exclusively decreased upon Ash1 RNAi. K36me2 was decreased reproducibly only in NSD RNAi while an Ash1-dependent decrease was observed in only one replicate (Supplementary Figure S2F). K36me3 was reduced in a Set2-dependent manner. The immunofluorescence staining did not reflect the reduction in K36me3 upon NSD RNAi found in Western blot, possibly due to the sensitivity limit on one hand and to the local distribution of the changes on the other hand (see below).

These results suggest that K36me1/2 do not simply serve as intermediates of trimethylation. They also reveal a complex interplay between the three HMTs, where each individual enzyme predominantly catalyzes one K36 methylation state, and only partially contributes to the others. Given the different distributions of K36me1/2/3 and HMT’s genome wide (Figure 1A-C), the differences may also be restricted to some genomic locations. We thus expect different phenotypes upon HMT loss of function mutants in flies. To explore this possibility, we reanalyzed published transcriptome data of NSD, Set2 and Ash1 knockout mutant fly brain (38), focusing on gene-level correlations. Indeed, the significantly deregulated genes poorly correlate between individual HMT mutants (Supplementary Figure S3), especially between Set2 and NSD, in support of the hypothesis of distinct roles of Set2 and NSD in gene regulation.

### Chromosome locus-specific H3K36 methylation changes upon HMT depletions

We next explored the genome-wide changes in K36 methylation after depletion of HMTs (Figure 3). Z-scores representing the direction and magnitude of change in K36me ChIP signal within 5-kb genomic bins were calculated for each HMT RNAi relative to the control (Figure 3A). To identify genomic regions that show significantly altered K36me1/2/3 upon RNAi, we adopted a statistical approach using the csaw pipeline (see Methods). The differentially marked regions obtained from this analysis were further visualized using high-resolution 2-kb chromoMaps to provide information regarding spatial patterns of methylation changes in each RNAi condition. For conciseness, only chromosome 3L is shown (all chromosomes are shown in Supplementary Figure S4). Profiles representing the respective K36me levels in unperturbed S2 cells above each chromoMap aid the interpretation of the relative changes while the gene annotation and H3K9me2 peak tracks illustrate local chromatin environment (Figure 3 B-D).

**Figure 3.**
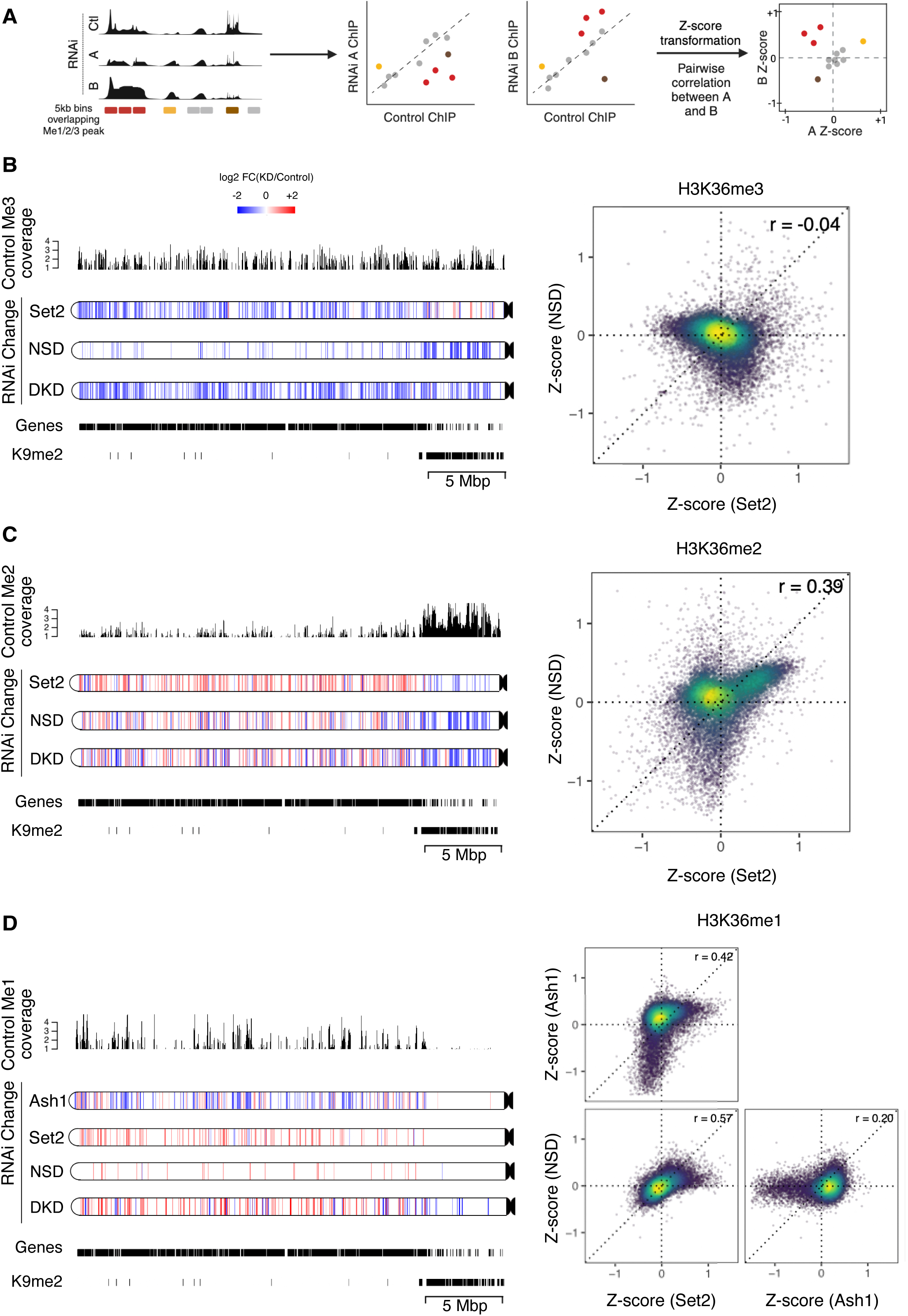
HMTs act on largely distinct genomic regions. A) Schematic depicting Z-score analysis workflow. Pairwise Z-score scatter plots representing correlations among changes in ChIP signal upon RNAi of HMTs with respect to control RNAi in genome-wide 5kb bins. Only bins overlapping at least one of K36me1/2/3 peaks in any RNAi condition were included. A negative Z-score indicates a reduction relative to control while a positive Z-score indicates an increase. B-D) Pairwise Z-score scatter plots for K36me1/2/3 upon RNAi of indicated factor (right). The color overlayed on the scatter indicates the local density of points. Pearson correlations are provided for each pair. Corresponding chromoMaps representing regions of significant difference in K36me3/2/1 signal for indicated RNAi conditions as derived from csaw analysis for chromosome 3L at 2-kbp resolution (left). The color of the regions (as indicated by the common scale in Figure 3B) represent log2-transformed value of number of normalized reads in RNAi condition relative to control condition. Control K36me3/2/1 signal overlayed above chromoMaps aid interpretation of relative changes. K9me2 peaks and gene annotations provided for reference.

The chromoMaps for K36me3 show that depletion of Set2 leads to a reduction of predominantly euchromatic K36me3, while depletion of NSD leads to loss of K36me3 at gene-poor pericentric heterochromatin and a subset of euchromatic bins (distinct from Set2-dependent regions as well as euchromatic K9me2 nanodomains) (Figure 3B). A few heterochromatic bins, which were marked by NSD-dependent trimethylation, were identified as significantly <u>increasing</u> in the Set2 knockdown condition.

To directly compare the differential effect of HMT depletion at higher resolution we visualized pairwise correlations at genomic bins that contain at least one K36 modification as Z-score scatter plots (see principle in Figure 3A). Off-center dots indicate the bins affected by Set2 or NSD or by both depletions while bins centered around (0,0) represent unchanging bins which are typically ‘background’ regions of low K36me3 signal (but are likely decorated by K36me1/2) (Figure 3B, C). K36me3 is reduced in a large fraction of genomic regions upon depletion of Set2 only (top-left quadrant), consistent with its role as a major trimethyl-transferase. Conversely, NSD RNAi leads to decrease of K36me3 in distinct genomic regions (bottom-right quadrant) which are not affected upon Set2 depletion, likely corresponding to PCH regions (and few euchromatic bins). Interestingly, depletion of either enzyme led to a mild gain of K36me3 at genomic regions methylated by the other enzyme (indicated by the positive shift from the horizontal or vertical axis, respectively), a phenomenon we term the ‘see-saw effect’. For example, in the absence of Set2 more K36me3 is found at pericentric heterochromatin, where NSD is strongly enriched (Figure 1). The see-saw effect can also be observed in several other instances. We assured that it was not due to a systematic bias in library preparation by verifying ChIP signals by qPCR at several diagnostic loci (not shown) and that Spike-in normalization does not change the results (see Methods). However, we emphasize that this mild reciprocal increase is often not identified as statistically significant by csaw, potentially due to limited detection power with three biological replicates. Nevertheless, this observation may suggest that the loss of one HMT somehow increases the activity of the remaining HMTs.

The contributions of NSD and Set2 to K36me2 are more complex (Figure 3C), most likely because K36me2 can arise from methylation (me0/1➔ me2) as well as from demethylation (me3 ➔ me2) reactions. The chromoMaps illustrate that K36me2 at heterochromatin and at certain euchromatic bins are dependent on NSD (compare NSD RNAi in Figure 3B, C) as also observed for K36me3 above. The corresponding bins where K36me2 is reduced in an NSD-dependent manner are in the bottom half of the scatterplot (Fig 3C). Conversely, K36me2 at heterochromatin was largely unaffected upon Set2 RNAi, although a mild decrease can be observed at a few bins which also display increased K36me3, hinting at the possibility that NSD is locally more active at converting K36me2 to K36me3 in the absence of Set2.

Further, an increase in K36me2 at many euchromatic bins can be observed upon NSD RNAi, despite unchanged K36me3, suggesting augmented K36me2-deposition by Set2. In contrast, the increased K36me2 levels upon depletion of Set2 are accompanied by corresponding drops in K36me3 levels and may thus be due to demethylation of K36me3 (compare Set2 RNAi in Figure 3B, C). These bins, which show increased K36me2 in both Set2 and NSD RNAi, are represented in the top right quadrant of the corresponding scatter plot (Figure 3C).

Lastly, chromoMaps and scatter plots highlight a strong loss of K36me1 upon depletion of Ash1 at loci with a strong enrichment of K36me1 in control cells confirming that Ash1 is the major K36me1-transferase in S2 cells (Figure 3D). Depletions of Set2 or NSD are rather associated with an increased K36me1 in distinct bins which have very little K36me1 in control cells, and likely result from demethylation of K36me2/3.

Taken together, our results suggest that the three methyltransferases establish the genomic K36 methylation signatures according to locus-specific rules. Individual HMTs may dominate the methylation state in certain spatial domains but may also indirectly affect each other’s activities exemplified by the see-saw effect which is significant at only few bins after csaw analysis.

### A gene-centric view of the K36 methylation landscape

The observation of distinct contributions of the three relevant HMTs to locus-specific methylation signatures prompted a more gene-centric analysis. K36 methylations are predominantly found on transcribed chromatin (Figure 1D). We selected around 10,500 genes enriched for at least one K36 methylation state in any one condition. The signals over the gene bodies were averaged and the resulting profiles clustered across all RNAi conditions. This yielded 12 clusters that once again illustrate the complexity of dynamic K36 methylation patterns across the genome. To facilitate a first interpretation of broad biological patterns while avoiding overinterpretation, we manually merged the clusters into three super-clusters (supercluster I = clusters 1-4, ochre; supercluster II = clusters 5-8, brown; super-cluster III = clusters 9-11, pink; Figure 4A). The minor supercluster IV (cluster 12, grey) contains a few hundred genes with very low K36me signals and was excluded from all subsequent analyses. The genes within the superclusters I to III are strongly correlated to specific K36 methylation profiles in steady state and tend to show consistent changes upon HMT depletion. Genome browser profiles for representative genes in each supercluster are shown in Figure 4B.

**Figure 4:**
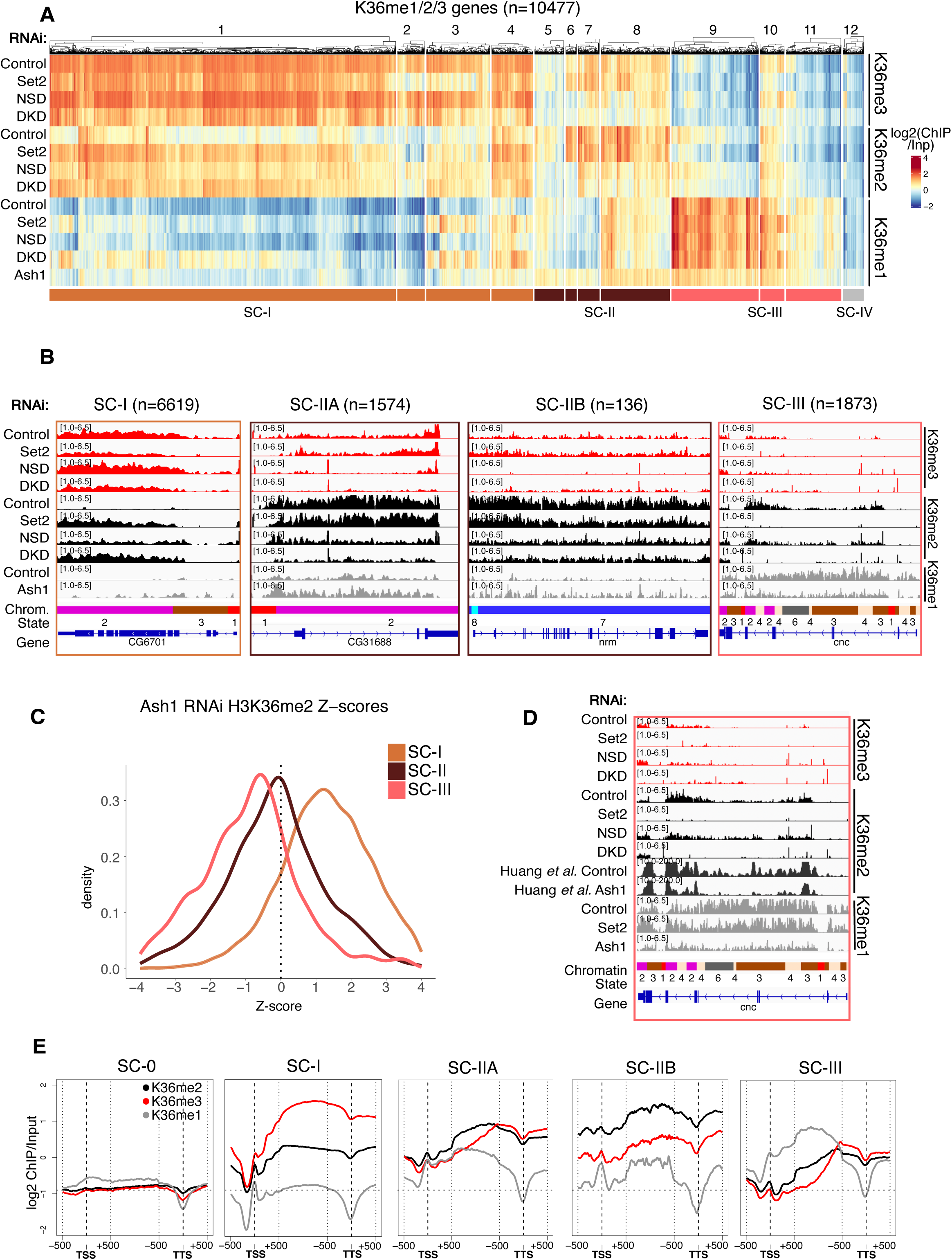
A gene-centric view of the K36 methylation landscape. A) Clustered heatmaps of gene body-averaged ChIP signal for K36me1/2/3 indicated on the right and RNAi condition indicated on the left. Only genes overlapping at least one of K36me1/2/3 peaks in any RNAi condition (n=10477) were used for clustering. Clusters are numbered 1-12 as indicated above the heatmap. The track below the main heatmap represents the manually grouping of clusters to define Superclusters (SC) which display similar patterns, resulting in ochre SC-I (n=6619), brown SC-II (n=1710) and pink SC-III (n=1873). B) Genome browser profiles of representative genes from each supercluster along with the number of grouped genes. Supercluster II was further classified into Euchromatic (IIA) or Heterochromatic (IIB) based on overlap with H3K9me2 peaks. RNAi condition and immunoprecipitation target indicated on the left; 9-state Chromatin States (as in Figure 1C) serve as reference. Representative genes shown for SC-I, IIA, IIB and III lie within clusters 1,8,7 and 9, respectively. C) Density plot of Z-scores representing change in K36me2 signal upon Ash1 RNAi (34) for superclusters defined in Figure 4A. D) Genome browser profiles of supercluster 3 representative gene highlighting effect of Ash1 RNAi on K36me2. RNAi condition and immunoprecipitation target indicated on the left; Chromatin States (as in Figure 1C) shown for reference. E) Cumulative plots for each cluster representing gene body relative distributions of K36me1/2/3. 1 kb regions centered around TSS and TTS are unscaled, while the rest of the gene body was scaled to 500 bins. SC-0 genes lack any detectable H3K36 methylation and serve as a reference for zero signal/baseline. This is also represented by the horizontal dotted line in all individual plots. Only genes of minimum length of 1500 bp were included in the analysis.

#### Supercluster I genes depend on Set2 for K36 methylation

The ochre supercluster I consists of genes which have the highest average K36me3 signal and very low me1/2 in control conditions (Figure 4B, C). At these genes K36me3 strictly depends on Set2. The reduction in K36me3 in absence of Set2 is accompanied by an increase in K36me2 and, most prominently in clusters 3 and 4, an increase in K36me1. Variable effects may be due to heterogeneity in locus-specific histone demethylase (HDM) activity (see below and Figure 6). We also observe a mild general increase in K36me2/3 upon depletion of NSD, reminiscent of the see-saw effect mentioned earlier and exemplified by the gene CG6701 (Figure 4A, B).

The combined depletion of both Set2 and NSD (DKD) results in patterns very similar to Set2 KD alone, although a some genes (particularly in cluster 4 and in cluster 1) show a slightly stronger reduction of K36me3 compared to Set2 knockdown alone. This might be due to mild contribution of NSD towards K36me3, which is not detectable in the individual NSD knockdown, because of compensatory Set2 activity (see-saw effect). Regardless, these observations show that Set2 is highly efficient in generating K36me3 with almost no observable intermediate K36me1/2 and mostly does not rely on prior methylation by NSD.

#### NSD is responsible for H3K36me2/3 at euchromatic and heterochromatic genes in supercluster II

Supercluster II comprises of 1710 genes with high K36me2, moderate K36me3, and variable K36me1 levels in unperturbed cells. Only a minor fraction of these genes (136) was located in pericentric heterochromatin and carried the heterochromatic K9me2 mark (Figure 4B, Supplementary Figure S5A). The abundant K36me2 is strongly reduced upon depletion of NSD, but unaffected upon depletion of Set2. The NSD-dependent reduction in K36me2 is in most cases accompanied by a reduction in K36me3, but no detectable changes for K36me1. Evidently, NSD is the major HMT responsible for K36me2/3 in this cluster of genes, of which 90% reside in euchromatin. For further downstream analyses (below), we distinguish euchromatic genes (supercluster SC-IIA) from heterochromatic NSD-dependent genes (supercluster SC-IIB) for clarity.

#### Ash1 is responsible for H3K36me1 in supercluster III

Supercluster III genes stand out because they are strongly marked by K36me1, which is markedly reduced by depletion of Ash1, but not affected by either Set2 or NSD depletion. This shows that K36me1 is a state in itself and not only a methylation intermediate (Figure 4A).

A subset of supercluster III genes are decorated by moderate K36me2 and low K36me3 (cluster 9) or moderate K36me2 and K36me3 (cluster 10). This K36me2/3-marking appears particularly Set2-dependent, suggesting that Set2 can add methyl groups processively to pre-methylated K36 (K36me1 and/or K36me2) in this particular context. Upon loss of Set2, these sites are demethylated, but K36me1 is maintained by Ash1. This response to Set2 depletion is markedly different from supercluster I where a widespread increase in K36me2 is observed. We used a previously published dataset (34) to show that K36me2 of genes within the supercluster III was indeed also dependent on Ash1 (Figure 4C, D). The simplest model explaining this co-dependence is that Set2 locally uses K36me1 placed by Ash1 to generate K36me2/3.

Remarkably, the above-mentioned see-saw effect is also visible upon Ash1 depletion on Supercluster-I genes which clearly gain K36me2 but not on supercluster-II genes (Figure 4C). However, a widespread (though mild) increase of K36me1 detected in supercluster II genes (Figure 4A, B) suggests that NSD-dependent *de novo* K36 methylation is increased in the absence of Ash1. Some Set2-dependent genes (in particular clusters 3 and 4) also experience a K36me1 increase upon Ash1 depletion. These genes probably overlap the K36me1-devoid bins that significantly gain K36me1 upon Ash1 RNAi in the previously shown chromoMap (Figure 3D) suggesting that Set2 and/or NSD contribute to K36me1 in certain contexts. Evidently, the see-saw effect is a rather locus-specific phenomenon (see discussion).

#### Context-dependent distribution of H3K36 methylation states along gene bodies

H3K36me3 is frequently referred to as a histone mark enriched towards 3’ ends of genes. Anecdotal genome browser observations (Figure 4B) suggest that the intragenic location of K36 methylation states differs. To investigate this more systematically, we generated cumulative profiles of K36 methylation states along scaled gene bodies between transcription start sites (TSS) and transcription termination sites (TTS), separately for each supercluster (Figure 4E). Unmethylated genes (SC-0) which were not included in the clustering serve as an internal negative control.

Genes in supercluster I show a strong 3’ bias for K36me3 but not K36me2, consistent with previous reports (66). K36me2 is broadly distributed throughout the bodies of the euchromatic genes in SC-IIA, whereas K36me1 drops towards the TTS and K36me3 increases sharply towards their 3’ ends. Such a modulation of K36me3 was not observed for the heterochromatic cluster II genes (SC-IIB), where the gene body signals are similar to those upstream of TSS and downstream of TTS. This observation fits the genome browser views (Supplementary Figure S1A) that show heterochromatic genes embedded in larger K36me2/3 domains.

Finally, at supercluster III genes, K36me1 broadly marks the gene bodies, whereas K36me2/3 are enriched towards the 3’ ends of transcription units. Notably, the intragenic distribution of the dimethyl mark is more similar to K36me3 rather than K36me1, further supporting the idea that most K36me2/3 at these genes is deposited by Set2 and not directly by Ash1 (Figure 4A).

The observation of distinct profiles of K36 methylation along transcription unit supports the concept of context-dependent methylation patterns.

### Genes of superclusters I, II and III are enriched in ePol, NSD and Ash1, respectively

The distinct distributions of K36 methylation marks at genes may be explained by targeting of the corresponding HMTs. If that was the case, we should find the HMT enriched at sites where K36 methylation is sensitive to its depletion. HMTs are often found enriched at distinct loci within larger domains of K36 methylation. To avoid averaging HMT ChIP coverage over the entire gene body, we selected the binding maxima for each HMT and calculated average signals within 2-kb windows around these maxima. The intensity distribution of the HMTs within the superclusters was visualized as violin plots (Figure 5A). A set of 3000 randomly sampled unmethylated genes (SC-0) served as an internal negative control.

**Figure 5:**
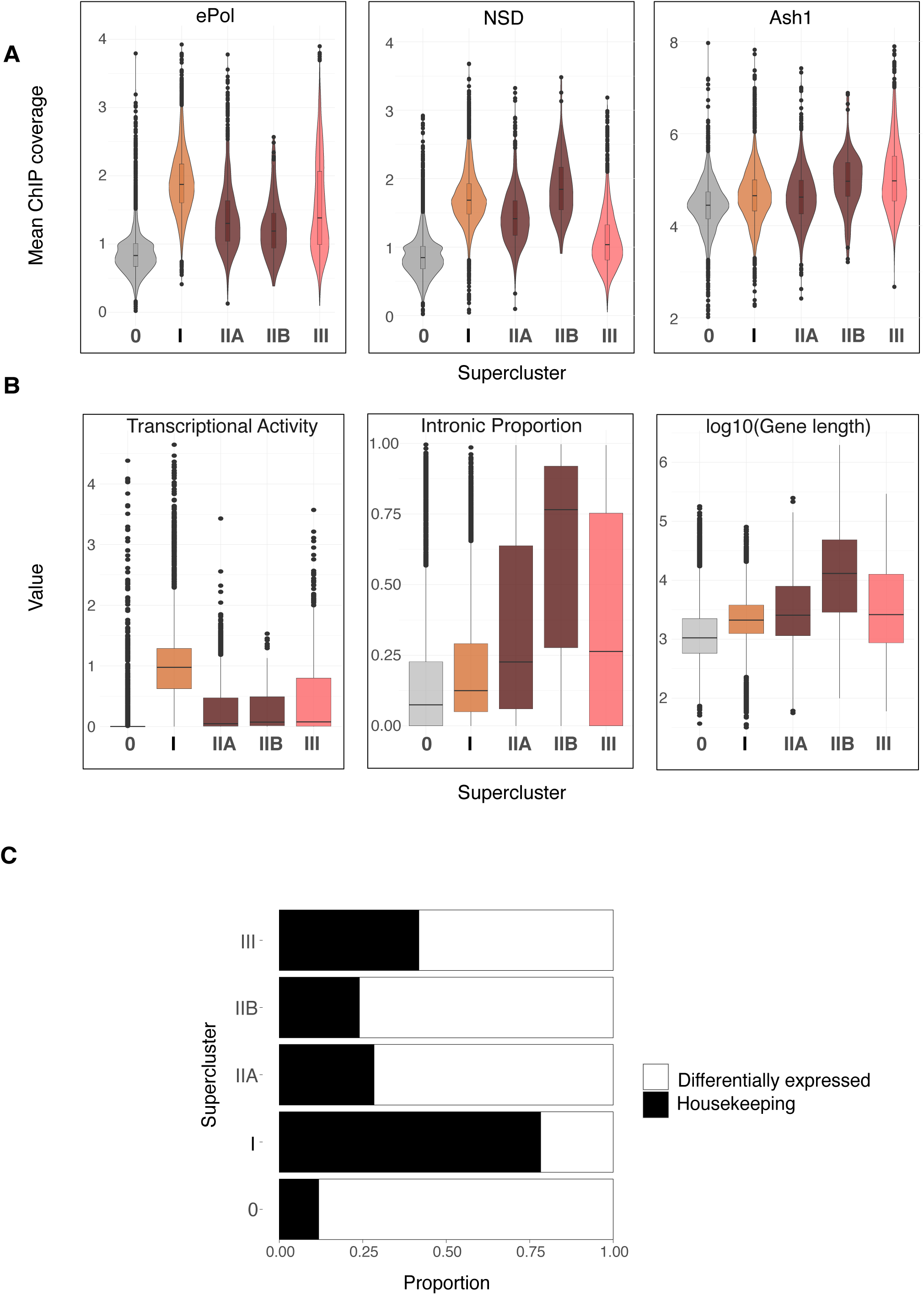
Gene clusters defined by HMT methylation patterns are correlated to different genic features. A) Violin plots representing average ChIP signal for 2 kb windows around HMT gene body peaks for NSD, ePol and Ash1 within superclusters defined in Figure 4A. Supercluster 0 represents randomly sampled genes (n=3000), which lack any detectable K36 methylation and serves as reference for zero signal/baseline. B) Boxplots showing the proportion of introns (sum of length of introns/total gene length), transcriptional activity (denoted by log10-transformed RNA-seq TPM values) and log10(gene length) for gene superclusters. C) Proportion of tissue-specific/invariant genes for gene superclusters based on FlyAtlas Expression Data for 25 tissues.

Set2 (defined by the presence of ePol) was strongly enriched on genes with Set2-dependent K36me2/3, most prominently supercluster I genes, followed by supercluster III genes (likely contributed by clusters 9 and 10) which show a Set2-dependent K36me2 and K36me2/3, respectively. Expectedly, NSD enrichment was stronger at heterochromatic genes (SC-IIB) compared to NSD-dependent euchromatic genes (SC-IIA). NSD was only weakly enriched at supercluster III genes. Set2-dependent supercluster I genes showed comparable levels of NSD to SC-IIA genes suggesting that NSD may be recruited to SC-I genes although its contribution to K36me at those genes is subtle. Lastly, analyzing the published Ash1 profile (34), we confirm that Ash1 was enriched the most in supercluster III. Mild enrichment within superclusters I and II-A may be due to overlapping domains (Figure 1A). The Ash1 enrichment observed within SC-IIB heterochromatic genes is probably contributed by non-specific signals which remain in Ash1 RNAi condition (data not shown).

Overall, these observations suggest that the three HMTs are the dominant drivers of K36 methylation at their highlighted gene clusters, but may also contribute to K36 methylation in other contexts.

### Housekeeping functions are marked by Set2, NSD and Ash1 target differentially expressed genes

The context for K36 methylation signatures (Figure 4E) may relate to gene-specific features like transcriptional activity, length and intron content. We visualized those features for genes in each supercluster as boxplots (Figure 5B). Genes in supercluster I were most highly transcribed and transcription activity correlated strongly with occupancy of the elongating RNA polymerase II (ePol) (Figure 5A, left). Genes in superclusters II and III are less expressed in agreement with lower ePol occupancy, tend to be longer and contain more introns. Long and intron-containing genes are often differentially expressed across cell types, in contrast to house-keeping genes.

To evaluate the degree of differential gene expression, we classified genes according to their FlyAtlas expression profiles for 25 different tissues (61), as either ‘tissue-invariant’ or ‘differentially expressed’ (Supplementary Figure S6A) and calculated the proportion of these two classes across the superclusters. This revealed that genes in superclusters II and III were predominantly differentially expressed, while supercluster I genes were ubiquitously transcribed (‘housekeeping genes’) (Figure 5C).

A further determinant of HMT targeting may relate to intron/exon content. K36me3 maps predominantly to exons, perhaps due to a slowing of ePol (and associated Set2) during splicing (4). Other studies also documented K36 methylation on introns (3). Anecdotal genome browser views (Figure 4B) hinted that K36me1/2/3 may mark introns and exons differentially in the different superclusters. We therefore examined the exon/intron distribution of steady state K36 methylation (Supplementary Figure S7A). We found K36me3 strongly enriched in exons versus introns in Set2-dependent supercluster I genes, whereas the bias is not observed for K36me2. The exon bias for K36me3 is substantial but also much less pronounced for K36me2 in the NSD-dependent SC-IIA genes and could be contributed by some genes that depend on both, Set2 and NSD (a certain fraction of cluster 7 genes). Interestingly, the bias is not lost upon Set2 depletion (Supplementary Figure S7B), suggesting that NSD may have a bias as well, despite no known interaction with ePol. Since a much weaker bias is observed for SC-IIB genes, exon selection is not an intrinsic feature of NSD but rather depends on the chromatin context (Supplementary Figure S7A).

Supplemental Figure S7B shows how the exon/intron distribution and bias shifts upon depletion of the HMTs and is revealing complex patterns that may be explored by future analyses.

The division of labor between HMTs for establishment of K36 methylation patterns genome-wide and the correlation to various genic features appears to be conserved during evolution. Similar SETD2-dependent and NSD1/2-dependent gene clusters were observed in two independent mouse data sets (Supplementary Figure S8).

### K36 methylation at transposable elements

Some transposable (TEs) are marked by H3K36me3 in *Drosophila* (29), and K36 HMTs were suggested to regulate TE expression (67). We mapped K36me2/3 at various repeat families across RNAi conditions and clustered the profiles (Supplementary Figure S9). Roughly 50% of TE families were not methylated in any condition. Among the families that were exclusively methylated by Set2, telomeric TE stand out, which are defined by high K36me3/low K36me2. Many transposon families that are characterized by NSD-dependent high K36me2 and moderate K36me3 levels are probably embedded in pericentric heterochromatin (68) as highlighted in the chromoMaps of Figure 3. These transposons also show increased K36me3/decreased K36me2 upon Set2 RNAi, probably resulting from the see-saw effect involving both NSD and Set2.

### H3K36me reader binding is determined by density and turnover of K36me3 and K36me2

Changes in K36 methylation patterns upon HMT depletion provide a unique opportunity to observe and compare the dependency of reader proteins on these marks. We explored the redistribution of exemplary reader proteins JASPer (29) and MSL3 (ChrX-specific) (21,28) upon HMT depletion for supercluster I and II genes, which comprise of almost all reader binding events (Figure 6A). The bulk levels of JASPer and MSL3 (inferred from other MSL complex proteins) were not affected upon Set2 or NSD depletion (Supplementary Figure S10A).

**Figure 6:**
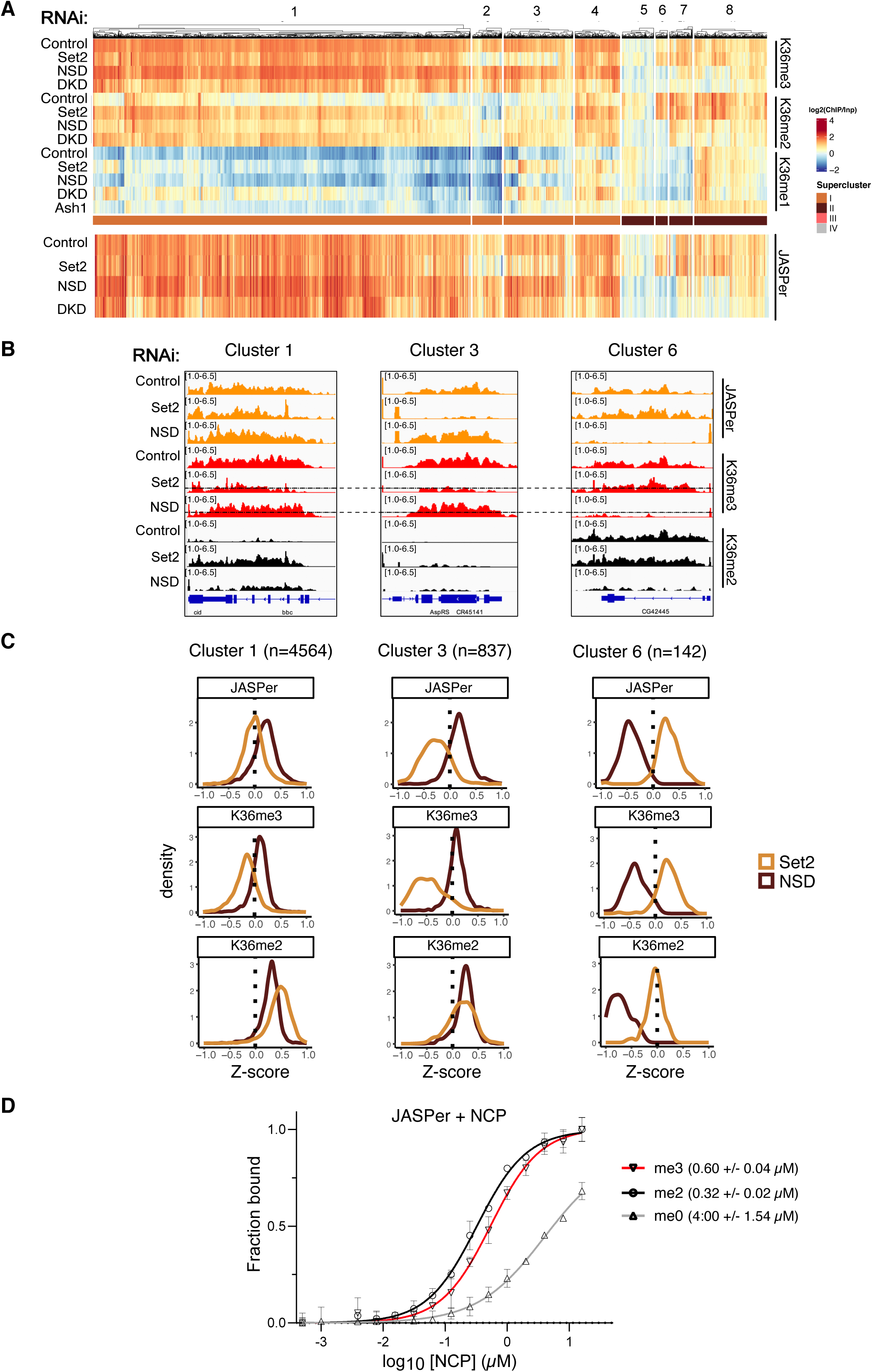
Robust binding of K36me3 readers above threshold K36me3 density. A) Heatmap of gene body average ChIP signal for JASPer in indicated RNAi condition ordered according to heatmap from Figure 4A. Note that supercluster III was excluded as it has very few genes bound by JASPer. B) Genome browser profiles for three distinct genomic loci highlighting differential response of JASPer to Set2 RNAi and NSD RNAi within indicated clusters. RNAi condition and immunoprecipitation target indicated on left-hand side. The dotted line in K36me3 ChIP in Set2/NSD RNAi represents the threshold derived from scatter plot in Supplementary Figure S10B. (Note that genome browser profiles display non-log transformed ChIP values. The calculated threshold of 1.35 corresponds to ~2.5 in non-log transformed units which is marked by the dotted line.) C) Z-score density plots representing direction and magnitude of change in gene body average ChIP signal for K36me2/3 along with reader JASPer for indicated representative clusters. D) Equilibrium binding between recombinant JASPer and unmodified, di- or trimethylated nucleosomes, respectively, determined using microscale thermophoresis (MST). Error bars represent the standard deviation from the mean values obtained from *n* = 2 experiments. Calculated dissociation constants are indicated within brackets.

Comparing the heat maps of JASPer binding upon HMT depletion (Figure 6A) revealed that JASPer follows K36me3 at some genes, in particular within clusters 2 and 3, but much less so in clusters 1 and 4. However, these latter sites are characterized by strong gains of K36me2 upon Set2 depletion. Upon depletion of NSD, JASPer binding is lost when both K36me2 and K36me3 are lost, most prominently in clusters 6, 7 and 8. Genomic profiles of representative genes in clusters 1, 3 and 6 illustrate the different scenarios inferred from the heat map (Figure 6B). These observations led us to hypothesize that JASPer may be attracted to both K36me3 and K36me2.

To explore this possibility further, we calculated Z-scores to quantify and compared the magnitude and direction of changes in reader binding and K36me2/3 signals upon HMT depletion in all genes of clusters 1, 3 and 6. At cluster 1 genes (Figure 6C, left panel), JASPer binding is unperturbed upon Set2 depletion. A significant reduction of K36me3 may be compensated by a corresponding strong increase of K36me2. In contrast, at cluster 3 genes, strong loss of K36me3 upon Set2 depletion is accompanied by only a mild gain of K36me2 and correlates with a loss of JASPer binding (Figure 6C, middle panel). In the third scenario, highlighted by cluster 6 genes, both K36me2/3 are eliminated in a coordinated fashion upon NSD knockdown resulting in reduction of JASPer binding (Figure 6C, right panels). These findings support the hypothesis that JASPer binds to both K36me2 and K36me3.

In order to generalize our observations to supercluster I and IIA genes, we visualized the change in JASPer binding (as Z-score) in response to Set2 depletion as a function of the absolute K36me3 ChIP signal (Supplementary Figure S10B). The scatter plot revealed that reduction in JASPer binding (Z<0) predominantly occurs when residual K36me3 levels fall below a threshold of 1.35. Below this threshold, decreasing K36me3 values result in a proportional JASPer reduction whereas JASPer continues to bind to ‘resistant’ genes that maintain higher levels of K36me3. Those resistant genes also acquire more K36me2 upon depletion of Set2, possibly because the turnover of both marks is linked (see below). Likewise, the same threshold of 1.35 is observed for the loss JASPer binding at SC-IIA genes upon NSD depletion (Supplementary Figure S10C). This 1.35 threshold is marked by a dotted line in all genome browser views (Figure 6B).

In light of the recent biochemical observation that the PWWP domain of LEDGF, the human ortholog of JASPer, can bind K36me3 and K36me2 with similar affinities (24), we measured the affinity of recombinant JASPer to K36me0/2/3 mononucleosomes by microscale thermophoresis (MST) (Figure 6D). JASPer was found to have a slight preference for K36me2 (K_D_=0.32 +/- 0.02 µM) over K36me3 mononucleosomes (K_D_=0.60 +/- 0.04 µM), whereas the affinity for unmodified nucleosomes was much lower (K_D_=4:00 +/- 1.54 µM).

We explored if the binding principles inferred from JASPer profiles also applied to the chromodomain protein MSL3. Because MSL3 binding is restricted to genes on the X chromosome, it cannot be displayed along with the reference heat map that clusters all genes. Nevertheless, representative X-chromosomal genes of the reference clusters 1, 3 and 6 show a similar behaviour and threshold effect for MSL3 (Supplementary Figure S10D, E). The corresponding density plots also show that MSL3 binding follows the general pattern of JASPer (Supplementary Figure S10F). At cluster 1 genes, MSL3 binding is unperturbed, whereas at clusters 3 and 6 MSL3 binding is lost along with K36 methylation in Set2 and NSD knockdowns, respectively. Because MSL3 binding profiles mirror that of JASPer, we hypothesize that MSL3 also binds to both K36me3 and K36me2. Altogether, MSL3 and JASPer leave chromatin when the combined density of K36me2 and K36me3 drop below a critical threshold.

The question remained why some genes in clusters 1 and 4 maintain relatively high levels of combined K36me2/3 despite significant depletion of Set2 during 7- or 10-day RNAi protocols (the duration of RNAi did not affect the general outcome, Supplementary Figure S11A), while genes in other clusters lose most K36me2/3 methylation. Conceivably, the resistance of K36 methylation at these loci upon HMT depletion is a function of turnover of the K36 methyl-marks, either due to K36 demethylation rates or turnover of K36me2/3-bearing nucleosomes, for example by transcription. In line with these considerations, we found more elongating RNA polymerase II (ePol) at cluster 2/3 genes that have much lower combined levels of K36me2/3 (‘sensitive genes’) compared to ‘resistant’ genes in clusters 1/4 (Supplementary Figure S11B). Furthermore, we found the putative K36 demethylase KDM2 (69), but not KDM4A, enriched at ‘sensitive’ clusters 2/3 genes. These findings suggest that robustness in K36me reader binding is determined by a complex interplay between dedicated HMTs and counteracting turnover mechanisms.

## Discussion

A variety of epigenetic processes have been linked to the methylation of histone H3 at lysine 36 in different organisms, mostly considering K36me2/3. Many of those processes influence the transcriptional output (5). Although K36 methylation is generally considered as an active mark, it is relevant for silencing by DNA methylation (13,14) and for silencing at facultative heterochromatin by H3K27me3 (15,16). In addition, K36me3 has been linked to genome stability (for review, see (70)) and HMTs establishing H3K36 methylation as well as K36me readers have been linked to cancer (71–73).

In mammals, HMTs that methylate K36 and corresponding reader proteins are numerous, suggesting partially redundant functions. To more easily uncover fundamental principles, we use the *Drosophila* model to explore the role of the three relevant HMTs Set2, NSD and Ash1 in the genome-wide distribution of the three K36 methylation states. We also determine the genomic locations of two dedicated readers proteins: the chromodomain MSL3, a subunit of the male-specific dosage compensation complex (DCC) (28) and the PWWP protein JASPer, which tethers JIL1 kinase to active chromatin, where it reinforces the active state by H3S10 phosphorylation (29). Importantly and in contrast to other methods, our optimized ChIP protocol enables improved mapping of K36 methylation and readers within constitutive heterochromatin (Supplemental Figure S1A), which are often poorly solubilized in traditional ChIP protocols (74,75) highlighting that they are not strictly active chromatin marks.

The mapping of each individual K36 methylation state in unperturbed conditions revealed complex patterns of distributions. Yet, the full complexity of the profiles only became apparent when the changes upon single or combined HMT depletion were scored in a gene-centric manner and the resulting heat maps subjected to unsupervised clustering. We hypothesize that the 12 clusters that emerged correspond to gene classes with shared properties and chromatin context. The grouping into superclusters served to outline broad features without getting lost in detail. The heterogeneity of the loss-of-function phenotypes revealed local, context-dependent activities of the HMTs.

### Ash1 establishes H3K36me1 domains overlapping enhancers

Monomethylation is the most abundant methylation state of K36 in *Drosophila* (65), but not well studied in other organisms. We found that K36me1 represents a defined chromatin state in regions with enhancers features, often in introns of differentially expressed genes. These sites cover about 15% of the genome and may correspond to the most densely K36-methylated domains, according to the relative abundance of the three methylation states (65). Remarkably, those sites bear very little K36me2/me3 (see exception below) and K36me1 appears often rather complementary to me2/me3 signals. Evidently, K36me1 is not only a methylation intermediate towards K36me2/3, but may have its own function at enhancers, which needs to be further explored. To our knowledge, readers that selectively recognize K36me1 are not known.

Most of K36me1 is placed by Ash1, an HMT that so far is mostly described as a K36 dimethylase important for restricting polycomb repression in flies and mammals (15,76). At other sites, such as a fraction of supercluster III genes, Set2 converts K36me1 placed by Ash1 into K36me2/3. The function of this locus- and context-specific methylation pathway at a large fraction of genic K36me1 enhancer domains remains to be explored.

### H3K36 methylation by NSD and Set2 represent largely independent pathways

Contrary to our expectations, in steady state conditions in S2 cells, NSD and Set2 act largely independently of each other at distinct chromatin domains and different genes. Set2 catalyzes K36me3 predominantly within intron-poor, highly transcribed housekeeping genes, while NSD catalyzes K36me2/3 in a heterochromatic environment as well as at numerous euchromatic, weakly transcribed genes with many introns and cell-specific expression profiles. Our data suggests that, in general, both NSD and Set2 can methylate nucleosomes *de novo* without prior ‘pioneering’ methylation by another enzyme. Set2 rarely adds a methyl group to K36me2 generated by NSD, in contrast to earlier interpretations of *in vitro* studies (30). Furthermore, we find that NSD is not only a K36 dimethylase but is also capable of generating K36me3 in *Drosophila*, which resonates with a recent report in mice (77). These results are in line with recent studies documenting distinct loss-of-function phenotypes of Set2 and NSD in flies (38,39). Moreover, these notions are corroborated by our reanalysis of independent mouse datasets, where we find that the mammalian K36 HMTs contribute to separate methylation profiles at distinct gene classes.

### Targeting and regulation of H3K36 methyltransferases

Unsupervised clustering of the K36 methylation changes at genes upon HMT depletion revealed a clear functional partitioning of the three HMT activities, which is, however, not exclusive. Each HMT appears to be most strongly enriched at loci, where depletion studies reveal their activity (Set2 in supercluster I genes, NSD at supercluster II and Ash1 at supercluster III). However, they can also be found at other sites, where they are apparently not active. For example, NSD is detected at supercluster I genes where K36 methylation largely depends on Set-2. Vice versa, Set2 can be mapped to supercluster II genes, where most K36me depends on NSD. These observations suggest that the functions of K36 HMTs can be regulated at two different steps, their targeting and the activation of enzymatic activity. This is reminiscent of yeast Set2, which is targeted to transcribed genes by interaction of the conserved SRI domain with the elongating RNA polymerase II. The SRI domain directly contributes to activation since its deletion abolishes K36me3 and reduces H3K36me2 activity *in vitro*. Further facilitation of set2 methylation activity is achieved in presence of monoubiquitinated Lys123 of H2B on nucleosome substrates (for review see (78)).

While Set2 is recruited to transcribed chromatin by interaction with the transcription machinery, it is less clear how NSD is targeted. It has been reported to be recruited to Beaf-32 bound active promoters (36), but we found no systematic association between Beaf-32 peaks and promoters of SC-II NSD-dependent genes (not shown). The molecular basis of NSD localization to heterochromatin was hypothesized to depend on HP1 (31), however preliminary experiments argue for an HP1-independent targeting mechanism (unpublished observations, and personal communication from D. Atkinson and T. Jenuwein). Sun *et al.* described a catalytic-activity independent role in promoting enhancer function by localizing to active enhancers (62). In contrast to these observations, we do not observe NSD at H3K27ac/H3K4me1 domains. For mammalian NSD enzymes, an autoinhibitory state is relieved by binding to nucleosomes (79) to trigger activation, NSD2 has been shown to be inhibited by H1 in nucleosomes (80). However, one conserved feature of all NSD orthologs is the presence of two PWWP domains, which could potentially modulate the activity of NSD and be implicated in a maintenance function. K36me3 is a relatively rare mark (65), thus it is also conceivable that NSD acts as a ‘maintenance’ methyltransferase by associating with a K36me2/3 mark. Such a scenario may serve to increase the local density of K36me3, to place K36me2 in the vicinity of K36me3 ‘seeds’ or to preserve the signature of active chromatin at poorly transcribed genes. Altogether, the targeting, activation and molecular function of *Drosophila* NSD, potentially through different mechanisms at euchromatin and heterochromatin, remains to be addressed in future studies. The function reported for mammalian NSD to establish faithful DNA methylation (13,81) will not be conserved in *Drosophila* which lacks DNA methylation-related machinery.

The determinants of genomic distribution of Ash1 have not been elucidated. Although we do not see extensive H3K36me2/3 dependent on Ash1 that is not dependent on Set2 it has been shown that *Drosophila* Ash1 in particular is activated by MRG15 (34,37).

### Indirect communication between HMTs: the see-saw effect

In several instances we observed that the depletion of one HMT led to an increase of the relevant mark at sites where another HMT is predominantly active, a phenomenon we termed see-saw effect. While the effect did not withstand rigorous statistical testing at our limited experimental power, it is nevertheless noteworthy. For example, Set2 depletion leads to an increase of K36me3 at pericentric heterochromatin, where NSD is strongly enriched. Similar effects can be observed across two different mouse data sets (13,62). Such an effect may be explained by competition for a shared substrate, such as SAM. Indirect communication between HMTs of this kind may lead to underestimation of co-dependencies between in particular Set2 and NSD in particular at genes where they colocalize if one activity is increased when the other one is depleted. However, it also highlights one possible mechanism that ensures robustness of H3K36me2/3 distribution upon metabolic perturbation. Interestingly, recent studies in mammals have linked H3K36me3 to the methionine cycle and vitamin B12 levels, which determine SAM availability, in reprogramming and cell differentiation (82,83).

### Robust readout of H3K36 methylation by readers bearing chromo- and PWWP-domains

At transcribed chromatin, the function of K36 methylation is mediated by dedicated reader proteins, of which we considered two, MSL3 and JASPer, which associate with methylated K36 with different domains. Both proteins behave similarly in our experiments. For MSL3, our systematic analysis extends previous studies (28,35), which either did not involve high-throughput sequencing or utilized Set2-deficient fly larvae where K36me3 is completely absent. Our RNAi approach provides a spectrum of intermediate K36me2/3 levels, which we exploit to obtain quantitative insights into reader binding.

We show that JASPer binds nucleosomes with K36me2 and K36me3 with similar affinity, as described for its mammalian ortholog LEDGF (24). This may explain why JASPer stays associated with genes that significantly lose K36me3, but retain K36me2. The ability of readers to recognize both states may ensure that transcribed chromatin is reliably bound upon changes. Furthermore, the fact that K36me3 is enriched at exons, while K36me2 is more prevalent at introns may assure that all aspects of transcribed chromatin are equally well addressed.

It appears that the chromatin interactions of both, the chromodomain reader MSL3 and the PWWP reader JASPer, depend on critical concentrations of K36me3/K36me2 regardless of which enzyme catalyzes the methylation. One way of rationalizing the observation is that the combined capacity of the K36me3/2 marks normally exceeds the requirements for productive tethering of the reader proteins, conferring robustness in reader binding against fluctuations during local turnover of histone or methyl-mark. While we formally determined a threshold value for K36me3, we note that this value represents a proxy for all potentially covarying chromatin modifications (exemplified by K36me2), which may contribute to reader binding in our model system. Loci where reader binding appears sensitive to Set2 or NSD depletion may have a higher turnover of H3K36me2/3 and/or a less efficient maintenance of K36 methylation. Corresponding physiological conditions may include changes in metabolic conditions (methionine or vitamin B12 availability, see above) for Set2-dependent house-keeping genes, and expression changes occurring during differentiation processes for NSD-dependent differentially expressed genes.

### Methylation read-out at heterochromatin

We found that K36me3 partitions between eu- and heterochromatin. The presence of K36me3, a mark globally associated with active chromatin, at H3K9-methylated heterochromatin has also been observed in other studies (3,77) and been described in human datasets (84,85). Such overlap of active with more repressive chromatin features may not simply be due to population heterogeneity since engineered dual readers validated the existence of such dual domains (77,86) and have been proposed to bookmark poised enhancers genome-wide in mouse (77). In *Drosophila*, the co-occurrence of those marks is mostly observed at genes embedded at pericentric heterochromatin, which also recruit JASPer and MSL3 readers. However, the relationship between K36me3/2 at PCH and reader binding is complicated as we also observe many instances of heterochromatic K36me2 domains which fail to recruit JASPer. Another heterochromatin modification/component may hinder binding to K36me2 locally. We thus speculate that the selective recruitment to K36me2/3 *in vivo* may be modulated in the chromatin context.

### Limitations of our study

Our systematic and comprehensive approach necessitated to focus on a simple cellular model. The fundamental principles we uncovered are likely to be modulated in cell-specific ways in the fly organism, where the expression of HMTs and KDMs vary spatiotemporally. For instance, NSD is maternally deposited and present during early development while Set2 contribution starts at NC5 with the minor zygotic gene activation.

Due to a lack of ChIP-grade antibody, we did not directly map Set2, but infer the genomic Set2 localization from the profiles of elongating RNA pol II.

Our antibody-based approach is blind to the H3 variant that bears the K36me mark and it is likely that histone H3.3 contributes to the observed ChIP signals. H3.3 is enriched at active chromatin, transposons and the hyperactive male X chromosome in *Drosophila* (87). Since H3.3 accounts for about 25% of all H3 and is placed where nucleosomes turn over, it is possible that a large fraction of the K36me marks we score reside on the variant (88).

The genomic annotation of introns and exons are based on whole-fly RNA datasets. It is possible that the long introns methylated by NSD might contain unannotated tissue-specific exons. Alternatively, K36me2/3 at these long introns may mark cell-specific regulatory domains (e.g., enhancers).

## Acknowledgements

We thank A. Campos-Sparr for technical assistance, S. Krause for conducting pilot experiments and R. Shetty for setting up CUT&RUN for H3K36me3. We are grateful to C. Wirbelauer and D. Schübeler from the FMI in Basel for sharing their HMT antibodies and A. Thomae from the Bioimaging Unit of the Biomedical Center, LMU for advice on microscopy. We thank T. Straub and T. Schauer from the BMC Bioinformatics Unit for access to the computational cluster, advice on bioinformatics and helpful scripts. We thank S. Krebs of LMU LAFUGA for Illumina sequencing. We thank V. Lux and E. Koutná from IOCB Prague for their help with the recombinant protein production and nucleosome reconstitutions. We thank the members of the Becker lab for sharing reagents, general advice and for comments on the manuscript.

## Data and code availability

The raw sequencing files in fastq format and the summarized genome browser tracks in bigwig format are available in the GEO database GSE253391 (private until accepted for publication). Custom code for analysis of data is available on Zenodo (https://doi.org/10.5281/zenodo.10474698). Western blots and Immunofluorescence images used for quantifications in Figure 2 and Supplementary Figure 2 are available on Zenodo (https://doi.org/10.5281/zenodo.10514405).

## Funding

MJ is member of the International Max-Planck Research School ‘Molecules of Life’ and the Integrated Research Training Group of the DFG-funded Collaborative Research Center ‘Chromatin Dynamics’. The research was funded by the German Research Council (DFG) through grants BE1140/11-1 to PBB and RE4742/1-1 to CR.

## Author Contributions

MJ, CR and PBB conceived the study. MJ carried out all experiments and performed the data analysis, except for MST experiments. MH purified recombinant JASPer, assembled modified nucleosomes and obtained MST measurements supervised by VV. CR established collaboration with VV. MJ, CR and PBB interpreted the data and wrote the manuscript with input from all authors.

## Declaration of Interests

The authors declare no competing interest.

---

**Supplementary Figure S1:**
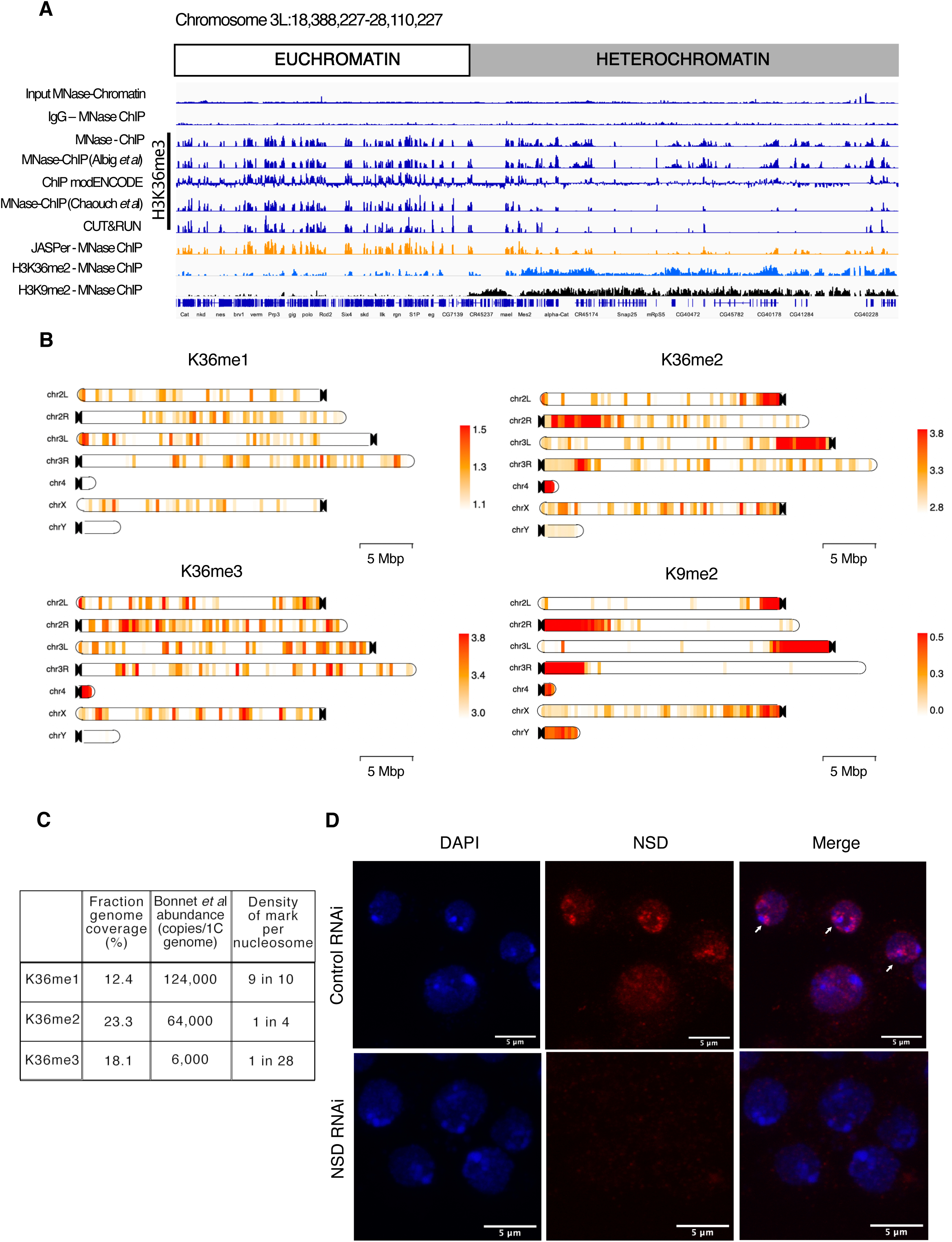
K36 modifications and HMTs are differentially distributed in male *Drosophila* cells. A) Genome browser profiles highlighting euchromatin bias introduced by the K36me3 mapping method. Comparison of different ChIP methods: MNase-ChIP: this work and Albig et al., (29) (includes mild shearing); CUT&RUN: this work; MNase-ChIP Chaouch et al., (58): native MNase-ChIP without crosslinking or shearing; ChIP modENCODE: Bioruptor shearing. IgG ChIP as well as chromatin input serve as negative controls. Colocalization of K36me3 signal with reader JASPer (MNase-ChIP) confirms specific signal. K36me2 coverage is provided for reference. H3K9me2 ChIP coverage demarcates heterochromatin and euchromatin. B) ChromoMaps representing steady-state enrichment of K36me1/2/3 in 10-kbp genomic bins for all *Drosophila* chromosomes. The H3K9me2 chromoMap serves as a reference for mappable pericentric heterochromatin domains. Note that continuous ChIP coverage is represented instead of peaks (representing regions of strongest signal) used in Figure 1B. C) Genomic coverage of K36me1/2/3 peaks and estimated densities based on Bonnet et al., (65). It should be noted that the density of K36me2 at heterochromatin is likely much higher than euchromatin. D) Immunofluorescence confocal microscopy of S2 cells treated with dsRNA against either GST (control) or NSD RNAi. Staining for DNA, NSD and merged images are shown. Arrowheads indicate chromocenter-proximal speckles of NSD. Scale bar is 5 µm.

**Supplementary Figure S2:**
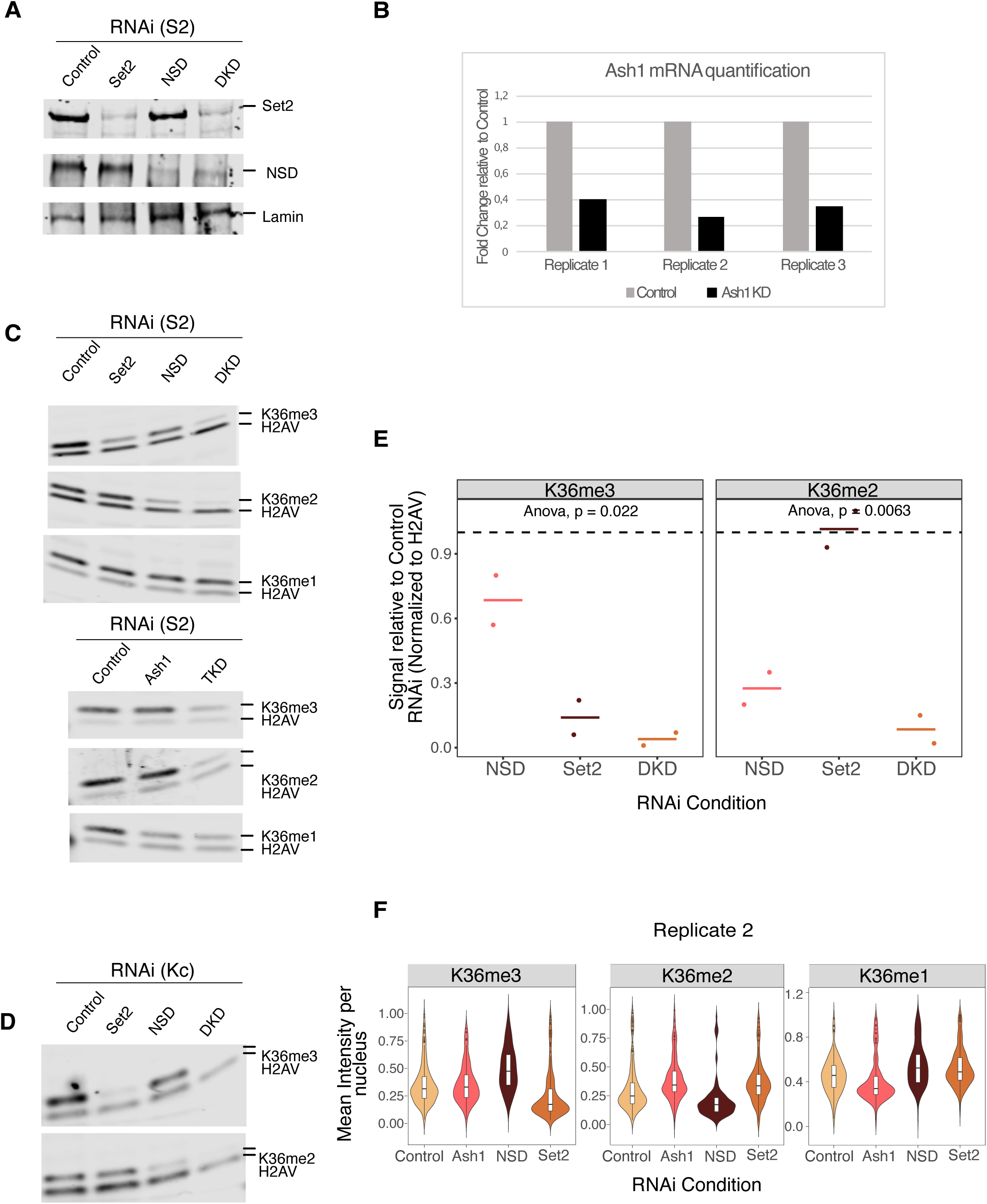
HMT depletions lead to similar alteration of bulk K36 methylation in male and female cells. A) Representative Western blot documenting the depletion of indicated HMTs in whole cell extracts from S2 cells treated with corresponding dsRNA. α-Lamin served as loading control. DKD: double knock down of Set2+NSD. B) RT-qPCR quantification of mRNA extracted from S2 cells treated with Ash1 dsRNA (N=3). C) Representative Western blots of K36 methylation marks used for quantification in Figure 2A. D) Representative Western blots of K36 methylation marks from Kc cells treated with GST (Control), NSD, Set2 or NSD+Set2 (DKD) dsRNA. E) Quantification of Western Blot analysis from Supplementary Figure S2D (N=2). F) Quantification of immunofluorescence images (n=~500 nuclei) from 2^nd^ biological replicate. First replicate presented in Figure 2C.

**Supplementary Figure S3:**
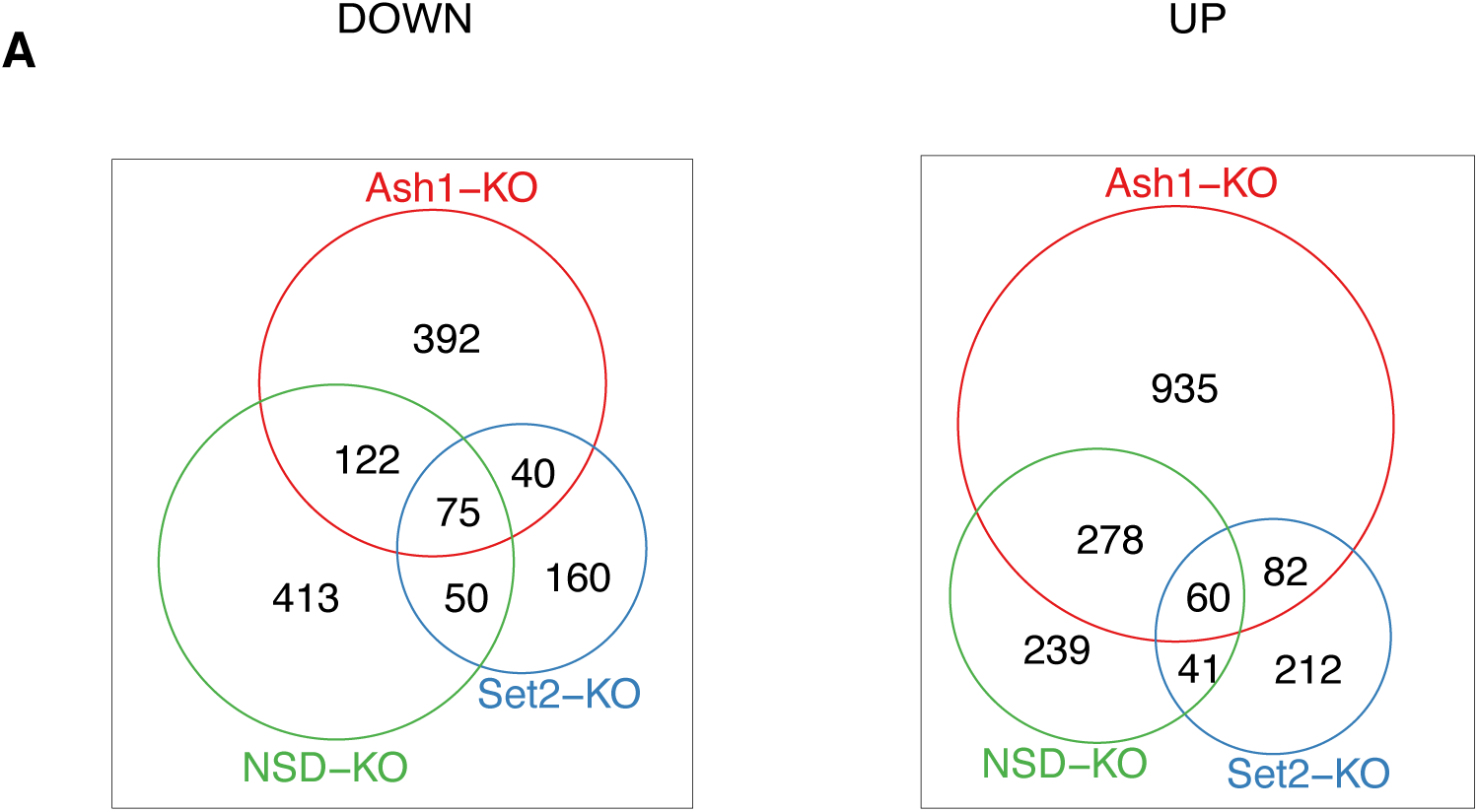
Poor overlap in transcriptomes of HMT Knockout *Drosophila* larval brains. A) Venn diagram representing overlap between significantly up-/downregulated genes (p<0.05, log2FC < −0.32 or log2FC > 0.26 (i.e. increased or decreased by at least 20%) from HMT knock-out (KO) Fly Larval Brain RNAseq data (38).

**Supplementary Figure S4:**
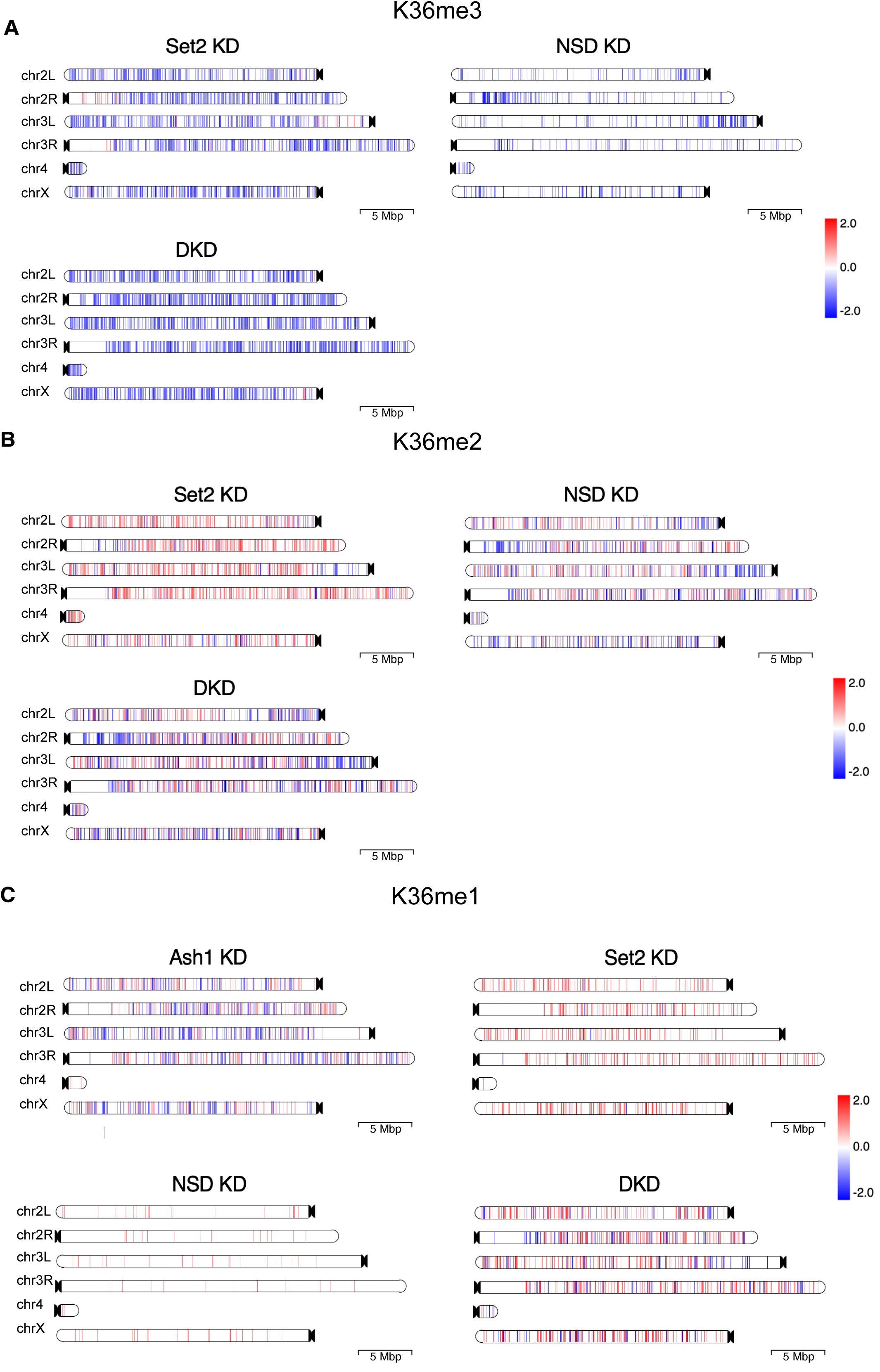
HMT Knockdown affects K36me1/2/3 in predominantly distinct genomic compartments. A-C) K36me3/2/1 chromoMaps representing regions of significant difference as derived from csaw analysis at 2-kbp resolution for all *Drosophila* chromosomes across all tested RNAi conditions. The color of the regions (as indicated by the common scale in Figure 3B) represent log2-transformed value of number of normalized reads in RNAi condition relative to control condition.

**Supplementary Figure S5:**
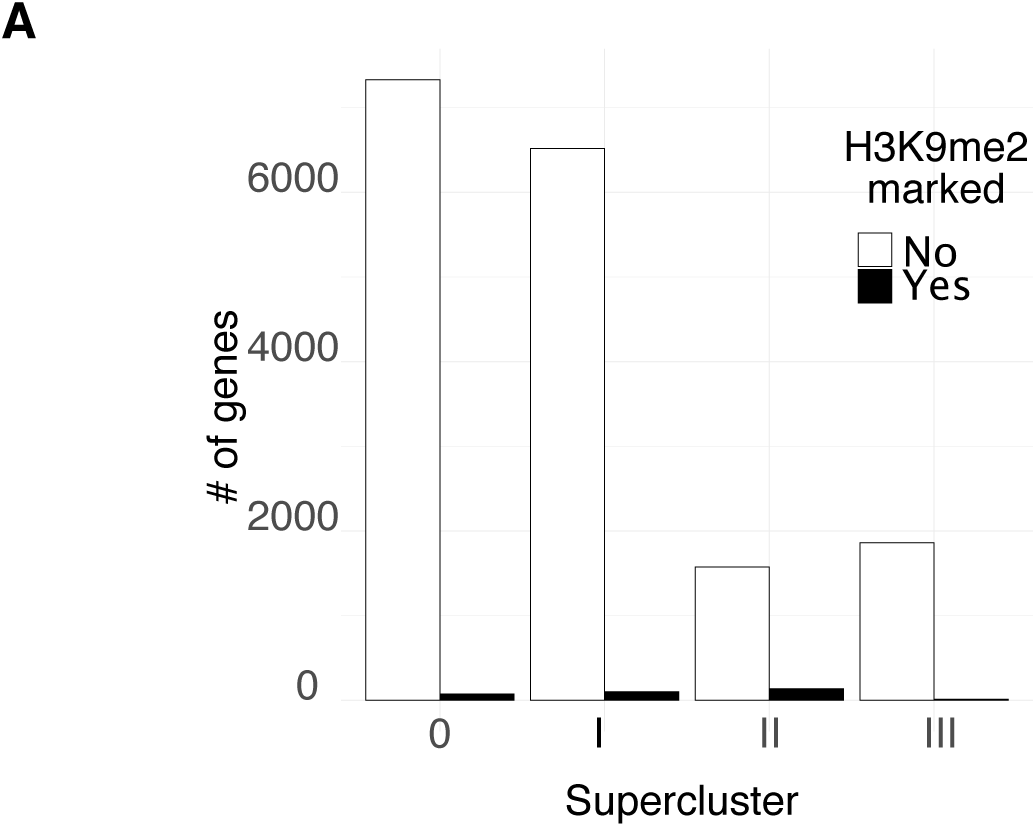
NSD-dependent methylation is present on many euchromatic genes. A) Quantification of proportion of Eu-/Heterochromatic genes for each supercluster-specific genes based on overlap with K9me2 peaks.

**Supplementary Figure S6:**
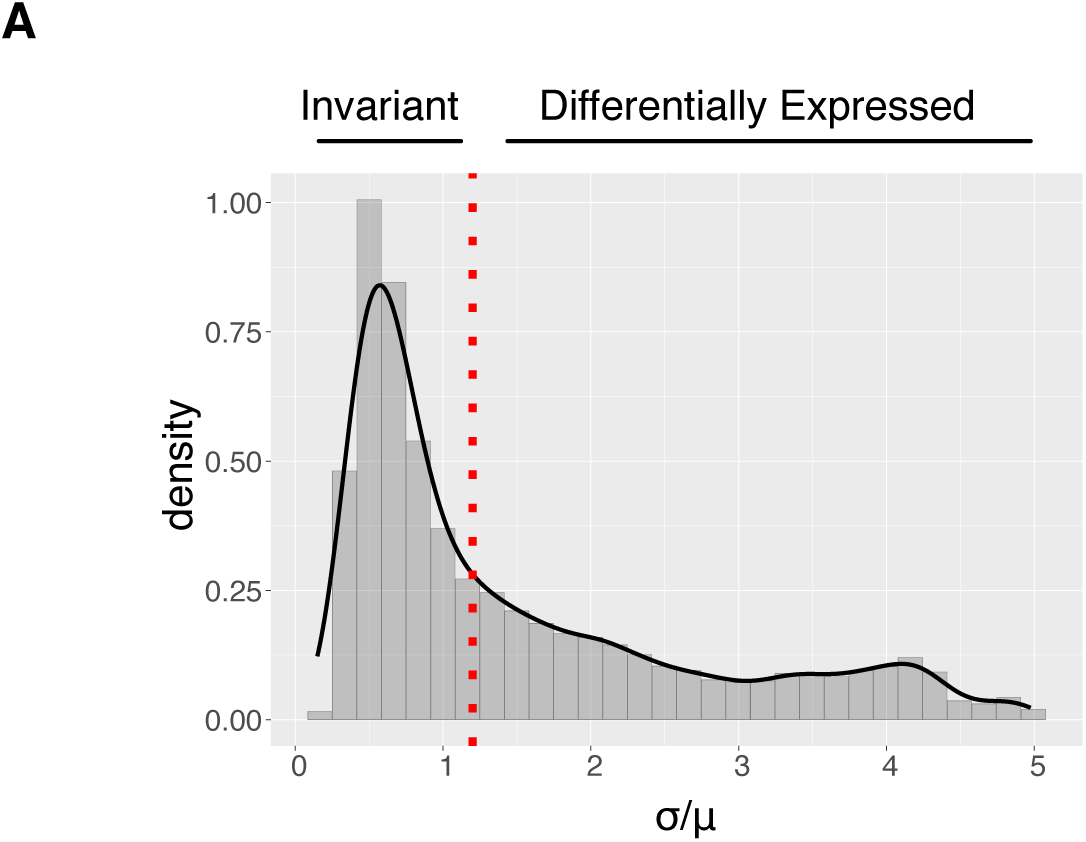
Classification of housekeeping (tissue invariant) and developmental (differentially expressed) genes based on FlyAtlas. A) Distribution of σ/µ values (σ = Expression Standard Deviation, µ = Expression Mean; calculated across 25 FlyAtlas Tissues). The red line represents the arbitrary threshold distinguishing tissue invariant and differentially expressed genes. Of note, genes with very low expression in all 25 tissues were automatically classified as differentially expressed as they are likely expressed in only a highly specialized cell type.

**Supplementary Figure S7:**
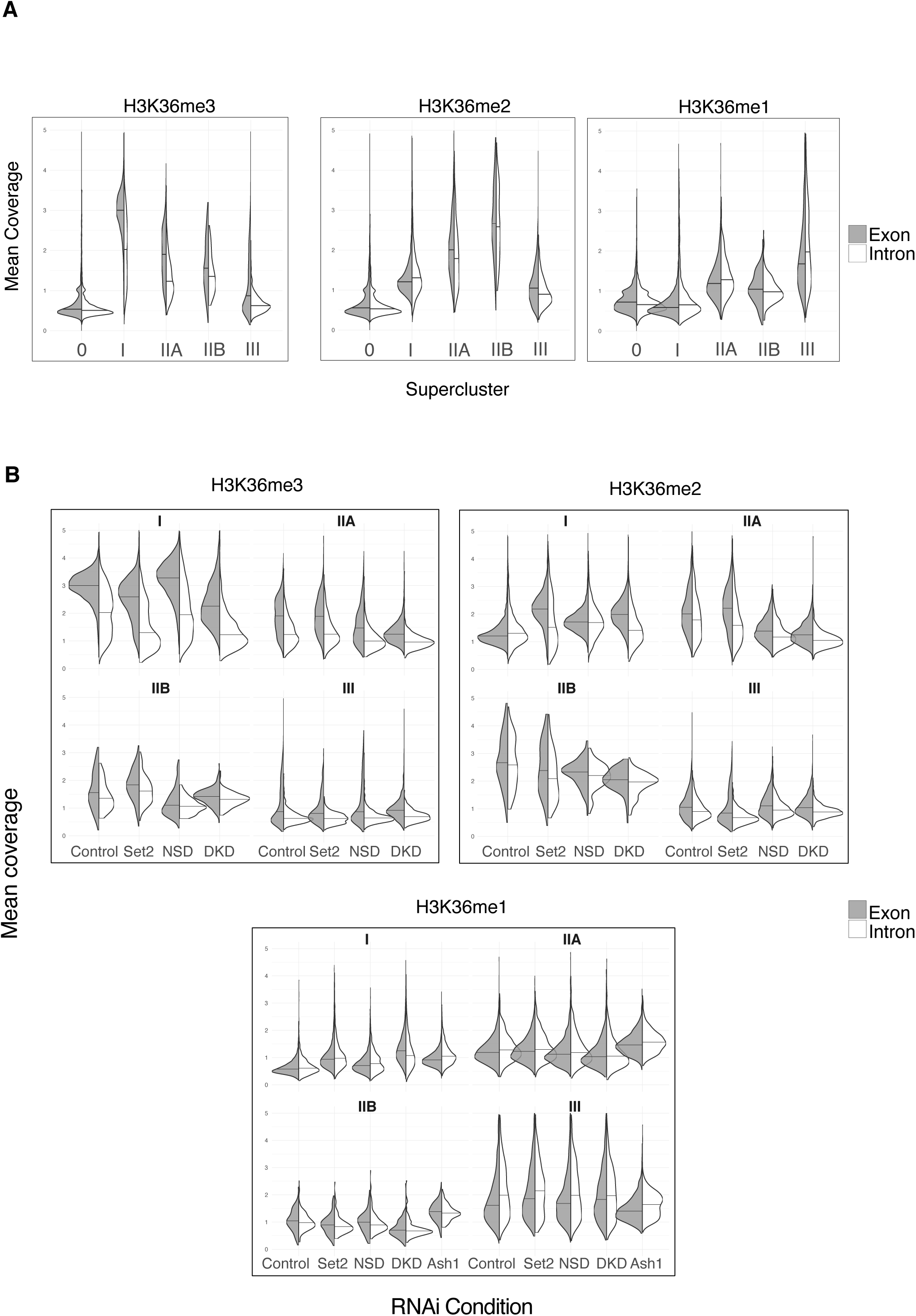
K36 methyltransferases show distinct preferences towards exons and introns. A) Split violin plots representing distribution of H3K36me3/2/1 ChIP signal in introns (white) and exons (gray) of genes in Control RNAi within each supercluster. B) Split violin plots representing changes in distribution of H3K36me3/2/1 ChIP signal in introns (white) and exons (gray) of genes in Control and HMT RNAi within each supercluster.

**Supplementary Figure S8:**
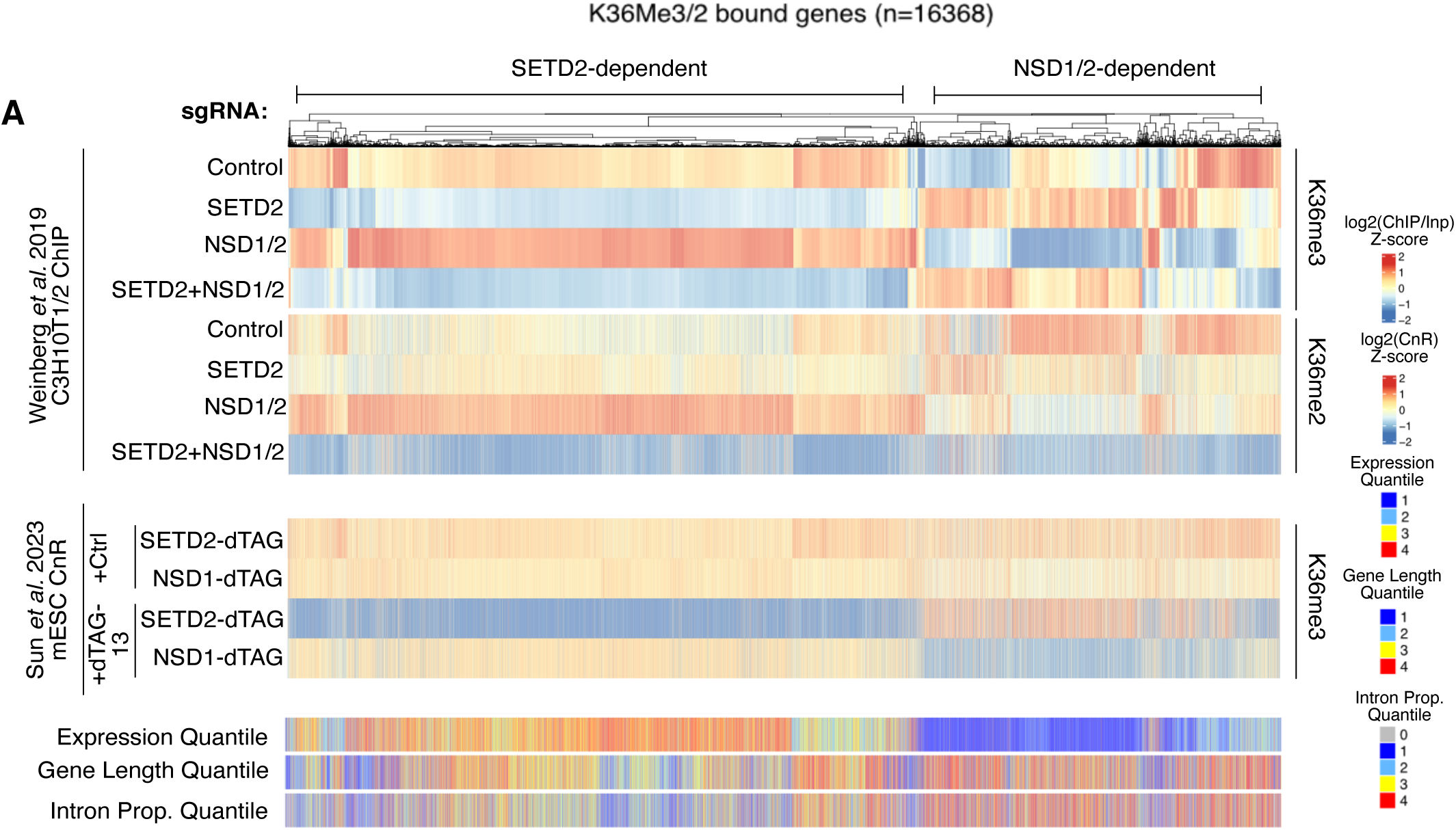
Set2/NSD-dependent methylation patterns are conserved in mouse cells. A) Top: Reanalysis of primary data from (13). Clustered heatmap representing gene body averaged ChIP signal for H3K36me3/2 (indicated on right) in Control or HMT Knockout background (indicated on left) mouse C3H10T1/2 cells. Only genes overlapping at least one of K36me2/3 peaks in Control cell line (n=16368) were used for clustering. Values were standardized for each gene (by subtracting mean and dividing by standard deviation) across KO conditions separately for K36me2/3 before clustering. Middle: Reanalysis of primary data from (62). Heatmap representing gene body averaged CUT&RUN signal for H3K36me3 in untreated or dTAG-13 treated HMT degron tagged mESC cells. Values were standardized across treatment conditions and are ordered based on the hierarchically clustered ChIP heatmap. Bottom: Heatmaps representing Quantiles of Expression, Gene Length and Intron Proportion ordered based on the ChIP heatmap (Quantile value 4=highest, 1=lowest; additional quantile category of 0 for Intron Prop. is to label genes with no introns). Expression data was obtained from Wang et al., 2019 The existence of NSD-dependent K36me3 domains was recently observed in mice (Barral et al., 2022). To verify if the genic methylation patterns that we observe are also conserved in mammals, we reanalyzed 2 additional recently published datasets (Weinberg et al., 2019; Sun et al., 2023). We first performed clustering analysis of genes under K36me2/3 peaks using H3K36me2/3 ChIP data from mouse C3H10T HMT Knockout (KO) cells (Weinberg et al., 2019) and correlated the clusters to H3K36me3 Cut&Run signal from mESC HMT-dTAG degron cells (Sun et al., 2023) as well as Gene Expression, Gene Length and Intronic Proportion Quantiles (Figure 5D). Of note, in this specific heatmap H3K36me2/3 signal was standardized (by subtracting the mean and dividing by standard deviation) across HMT perturbation conditions to robustly cluster genes which show similar patterns and thus doesn’t reflect absolute ChIP/CnR signal values. But overall, the absolute K36me3 signal at NSD1/2-dependent genes is about 3-10 fold lesser than SETD2-dependent genes depending on the mapping methodology used (as CnR strongly underestimates H3K36me3 signal at NSD-dependent genes). We clearly observe two distinct responses in the HMT KO/dTAG-13 treated cells where SetD2 deposits H3K36me3 independent of NSD1/2 at highly expressed, short, exonic genes while NSD1/2 catalyzes H3K36me2/3 at poorly expressed, long, intronic genes. Intriguingly, the “see-saw” effect described in *Drosophila* is also apparent in these datasets where NSD1/2 is more active upon SetD2 depletion. Of note, the TKO me2/3 patterns are not a simple addition of the individual knockouts, which may hint at the involvement of other mammalian H3K36 HMTs (like SETD5 and/or NSD3) (Sessa et al, 2023) or altered demethylases localization. Importantly, these patterns are clearly visible across two very different perturbation strategies (Long-term Knockout vs Acute Inducible Degradation) and mapping methodology (ChIP vs CUT&RUN), supporting robustness of the phenomenon.

**Supplementary Figure S9:**
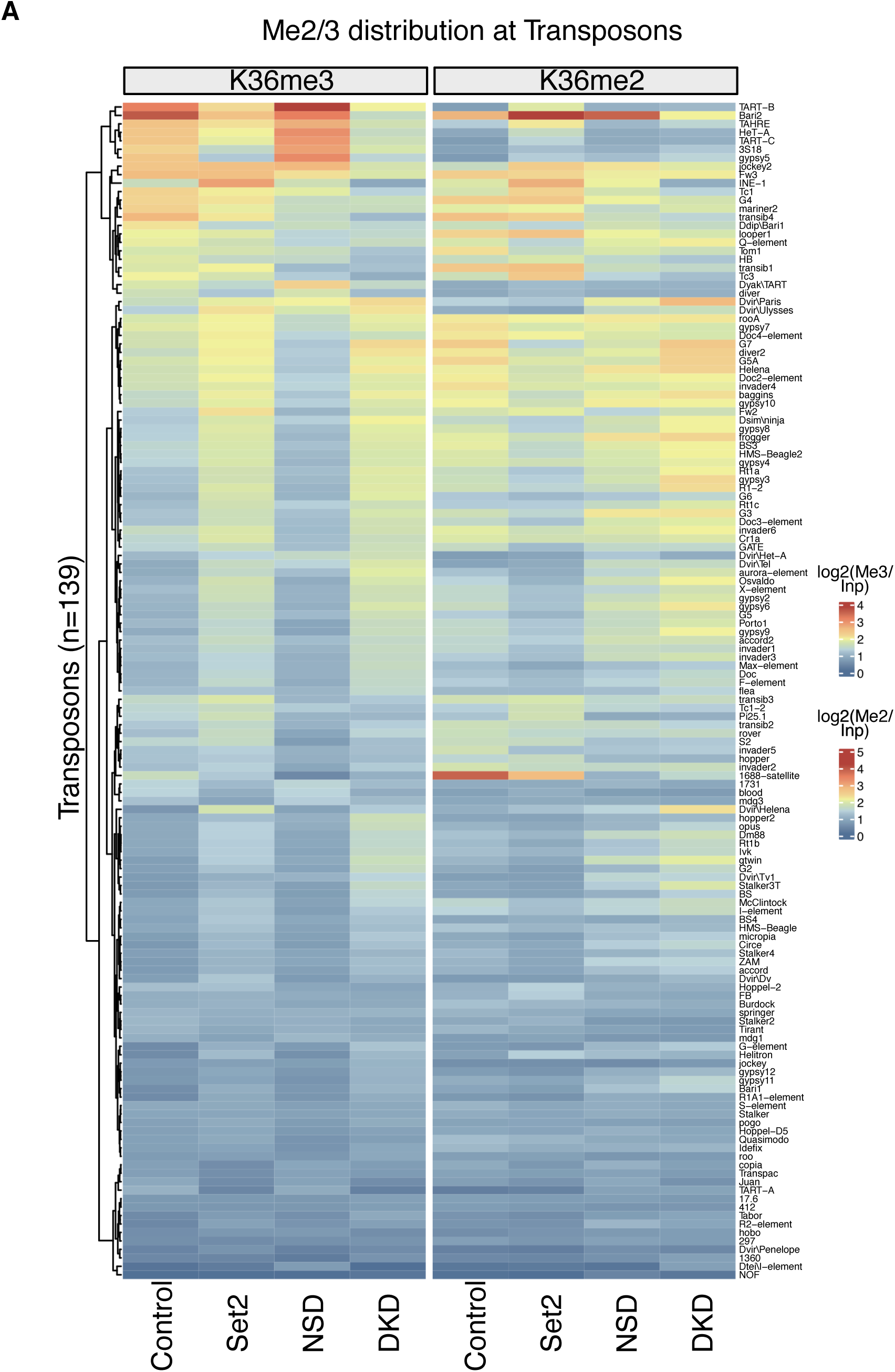
Set2 and NSD deposit K36me2/3 on distinct transposons. A) Clustered Heatmap representing average K36me2/3 ChIP enrichment at 139 transposon families (names indicated on the right) in GST (Control), Set2, NSD or Set2+NSD (DKD) RNAi.

**Supplementary Figure S10:**
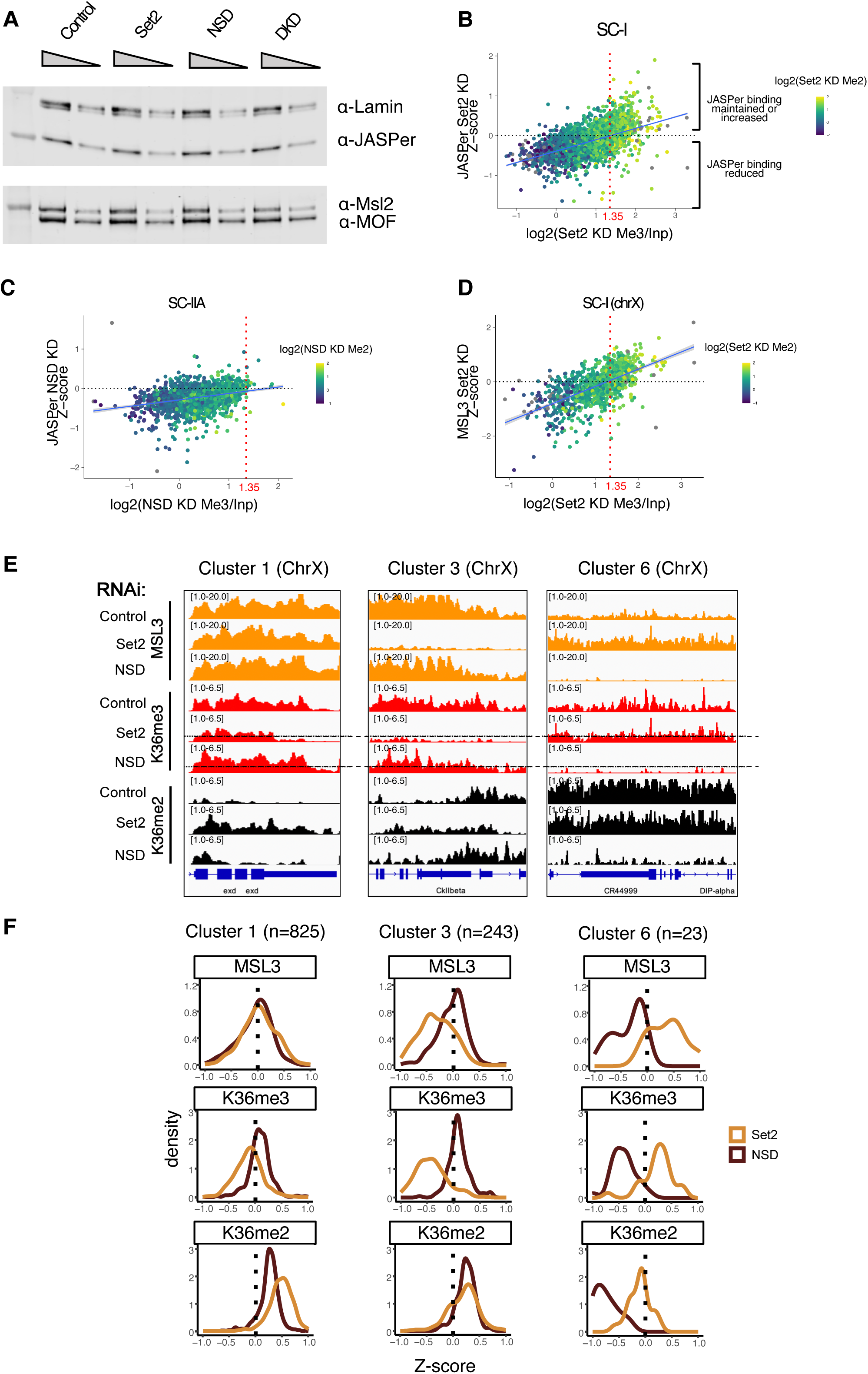
H3K36me reader binding is determined by threshold density of K36me2/3. A) Western blot analysis using α-JASPer, α-Lamin, α-Msl2 and α-MOF antibodies and S2 whole cell extracts treated with Control, Set2, NSD or Set2+NSD (Comb) RNAi. Two dilutions (1x and 0.5x were loaded per sample) B) Scatter plot representing JASPer Set2 RNAi Z-score (y-axis) as a function of K36me3 ChIP value in Set2 RNAi (log_2_ transformed) for SC-I genes. Each point represents a gene. The color of the points represents the K36me2 ChIP value in Set2 RNAi (log_2_ transformed). The red dashed line denotes the x-axis intercept of the best fit line and represents the threshold K36me3 value below which majority of the genes have a negative JASPer Z-score. C) Scatter plot representing JASPer NSD RNAi Z-score (y-axis) as a function of K36me3 ChIP value in NSD RNAi (log_2_ transformed) for SC-IIA genes. The color of the points represents the K36me2 ChIP value in NSD RNAi (log_2_ transformed). The red dashed line corresponds to the threshold derived from Supplementary Figure S10B. D) Scatter plot representing MSL3 Set2 RNAi Z-score (y-axis) as a function of K36me3 ChIP value in Set2 RNAi (log_2_ transformed) for SC-I ChrX genes. The color of the points represents the K36me2 ChIP value in Set2 RNAi (log_2_-transformed). The red dashed line corresponds to the threshold derived from Supplementary Figure S10B. E) Genome browser profiles for three distinct representative genes on the chromosome X highlighting differential response of MSL3 to Set2 RNAi and NSD RNAi within indicated clusters. RNAi condition and immunoprecipitation target indicated on left-hand side. The dotted line in K36me3 ChIP in Set2/NSD RNAi represents the threshold derived from scatter plot in Supplementary Figure S10B. F) Z-score density plots representing direction and magnitude of change in gene body average ChIP signal for K36me2/3 along with reader MSL3 for indicated representative clusters restricted to chromosome X.

**Supplementary Figure S11:**
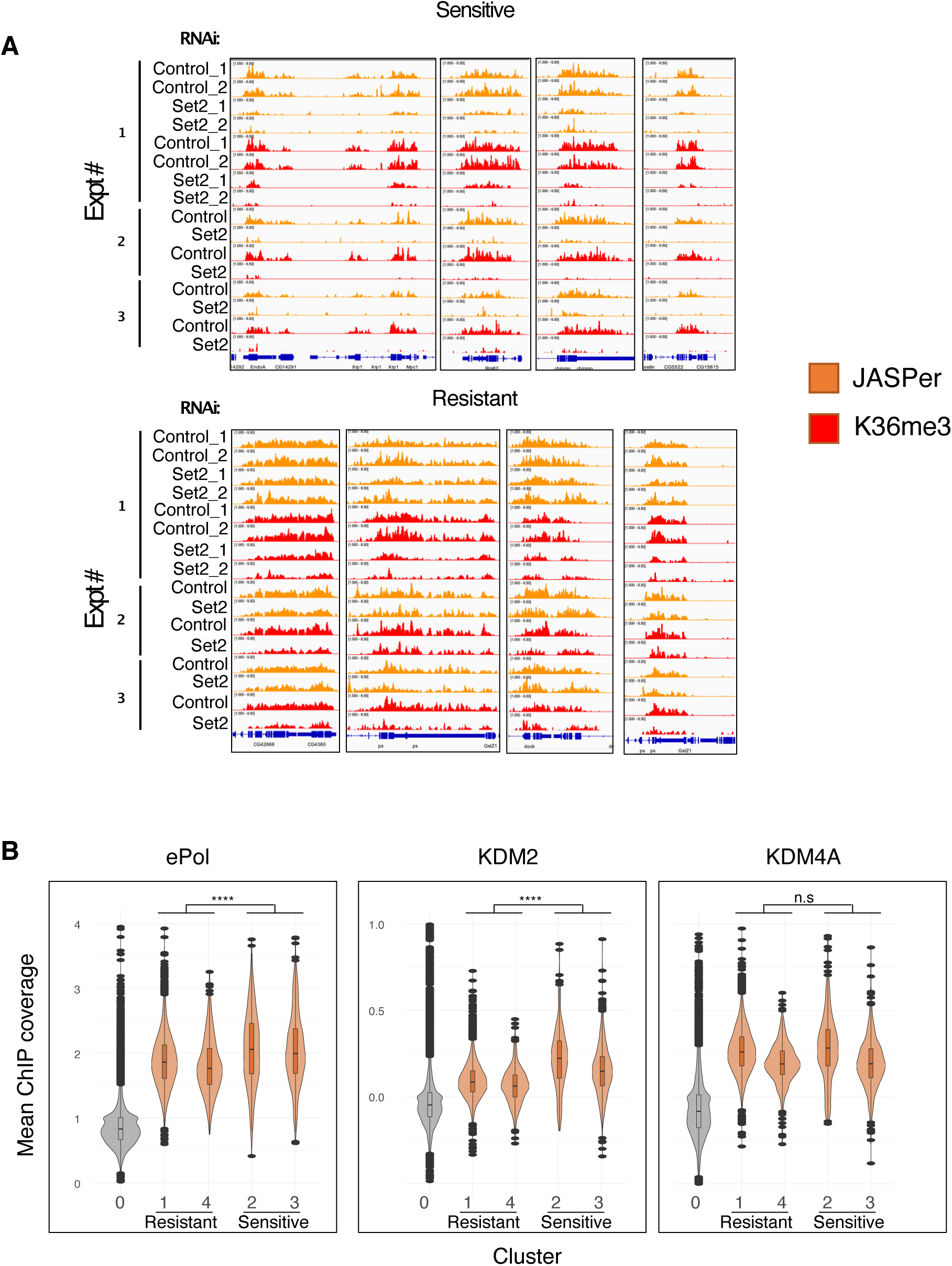
Factors involved in turnover of K36me2/3 may dictate ‘resistant’ or ‘sensitive’ outcomes upon Set2 RNAi. A) Genome browser profiles for multiple genomic regions documenting reproducible ‘sensitive’ and ‘resistant’ phenotypes of K36me3/JASPer across replicate experiments of 7 day RNAi (Expt #1) or 10 day RNAi (Expt #2,3). B) Violin plots representing gene-body average ChIP signal for HMT (ePol) and HDMs (KDM2 and KDM4A) for clusters within SC-I. Cluster 0 represents genes which lack any detectable H3K36 methylation and serves as a reference for zero signal/baseline. Definition of ‘sensitive and ‘resistant’ clusters follows from main text. One-sided t-test was performed to identify factors selectively enriched in the ‘Sensitive’ clusters relative to the ‘Resistant’ clusters (**** denotes a significance value of p < 0.001, n.s=Not Significant).

## Notes

### Competing Interest Statement

The authors have declared no competing interest.

